# GLUT4 dynamic subcellular localization is controlled by AMP kinase activation as revealed by proximal proteome mapping in human muscle cells

**DOI:** 10.1101/2023.06.06.543897

**Authors:** Anuttoma Ray, Jennifer Wen, Lucie Yammine, Jeff Culver, Jeonifer Garren, Liang Xue, Katherine Hales, Qing Xiang, Morris J. Birnbaum, Bei B. Zhang, Mara Monetti, T.E. McGraw

**Author notes:** <u>Corresponding author</u>.

## Abstract

Regulation of glucose transport into muscle and adipocytes, central for control of whole-body metabolism, is determined by the amount of GLUT4 glucose transporter in the plasma membrane (**PM**). Physiologic signals (activated insulin receptor or AMP kinase [**AMPK**]), acutely increase PM GLUT4 to enhance glucose uptake. Here we show in kinetic studies that intracellular GLUT4 is in equilibrium with the PM in unstimulated cultured human skeletal muscle cells, and that AMPK promotes GLUT4 redistribution to the PM by regulating both exocytosis and endocytosis. AMPK-stimulation of exocytosis requires Rab10 and Rab GTPase activating protein TBC1D4, requirements shared with insulin control of GLUT4 in adipocytes. Using APEX2 proximity mapping, we identify, at high-density and high-resolution, the GLUT4 proximal proteome, revealing GLUT4 traverses both PM proximal and distal compartments in unstimulated muscle cells. These data support intracellular retention of GLUT4 in unstimulated muscle cells by a dynamic mechanism dependent on the rates of internalization and recycling. AMPK promoted GLUT4 translocation to the PM involves redistribution of GLUT4 among the same compartments traversed in unstimulated cells, with a significant redistribution of GLUT4 from the PM distal Trans Golgi Network Golgi compartments. The comprehensive proximal protein mapping provides an integrated, whole cell accounting of GLUT4’s localization at a resolution of ∼20 nm, a structural framework for understanding the molecular mechanisms regulating GLUT4 trafficking downstream of different signaling inputs in physiologically relevant cell type and as such, sheds new light on novel key pathways and molecular components as potential therapeutic approaches to modulate muscle glucose uptake.

## Introduction

Muscle cells and adipocytes are the major sites of postprandial glucose disposal^1-3^. In both cell types, glucose uptake is low when blood insulin levels are low, and postprandial elevated insulin increases glucose uptake by inducing a redistribution of the GLUT4 glucose transporter from intracellular compartments to the plasma membrane (**PM**)^2^. GLUT4 redistribution to the PM is impaired in insulin resistance and type 2 diabetes, warranting a thorough understanding of the underlying molecular mechanisms governing GLUT4 translocation in muscle and fat cells. In skeletal muscle, contraction (and exercise) also stimulates glucose uptake, and this too is dependent upon GLUT4 translocation to the PM^1^. The molecular and mechanistic overlaps between insulin and contraction signaling inputs to GLUT4 are not well understood. At a functional level it is known that 1) a bout of exercise sensitizes muscle to a subsequent challenge with insulin, and 2) exercise-induced GLUT4 translocation is unaffected by insulin resistance, revealing similarities yet important differences in the control of GLUT4 trafficking downstream of these two physiologically relevant signaling inputs^4,5^.

AMP–activated protein kinase (**AMPK**) is a serine/threonine kinase that acts as a sensor of cellular energy status activated by increased [AMP]/[ATP] ratio^6^. AMPK activation in muscle induces glucose uptake downstream of GLUT4 translocation to the PM^7^, and AMPK activation is required for exercise to enhance subsequent muscle insulin-sensitivity^8,9^; observations that support AMPK as a possible therapeutic target controlling blood glucose^10^. Despite AMPK and insulin receptor signaling converging on GLUT4 trafficking, the interplay between these signal transduction mechanisms has yet to be described in complete molecular detail.

Two Rab GTPase activating proteins (**GAP**s), TBC1D1 and TBC1D4(AS160), have emerged as critical regulators of GLUT4 trafficking through their regulation of Rab proteins^2^. Rab10 has been shown in studies of cultured adipocytes and adipocyte-specific knockout mice to be downstream of TBC1D4 in insulin-stimulated GLUT4 translocation^11-14^. Whereas Rab8a has been shown in studies of rodent muscle cells to be required for insulin stimulated GLUT4 translocation^15^. The Rab(s) required for AMPK-stimulated GLUT4 translocation in muscle remain to be described, although electrical stimulation-induced GLUT4 translocation in cultured muscle cells requires Rab8a, Rab13 and Rab14^16^.

Here we use a human skeletal muscle cell line to investigate the molecular machinery governing GLUT4 trafficking in unstimulated cells and to explore how GLUT4 trafficking is altered by insulin receptor or AMPK activation. Insulin induces redistribution of GLUT4 to the PM by accelerating exocytosis, consistent with previous studies of rodent muscle cells^1^, whereas AMPK stimulation promotes GLUT4 translocation by regulating both exocytosis and endocytosis. Our kinetic studies support a model in which PM GLUT4 is in equilibrium with the PM in all conditions, with the amount of PM GLUT4 determined by the kinetics of GLUT4 trafficking to and from the PM. Because GLUT4 trafficking among distinct membrane compartments controls GLUT4-dependent glucose uptake, understanding this process requires a spatial description of GLUT4’s itineraries in different physiologic states. Spatial information is usually provided by imaging studies, in which colocalizations with established fiduciaries are used. However, imaging studies provide a "low density" map of GLUT4 localization because no more than a handful of fiduciaries can be used at any one time. Here we report on the high-density, high-resolution GLUT4 proximal proteome of human skeletal muscle cells generated by APEX2 biotinylation mapping. The GLUT4 protein atlas reveals the intracellular compartments traversed by GLUT4 in unstimulated cells and how GLUT4’s distribution among those compartments is altered by AMPK activation. These findings provide a framework for studies to define the molecular mechanisms regulating GLUT4 trafficking downstream of distinct signaling inputs relevant to metabolic disease states in a physiologically relevant cell type. The new insights on the regulation and the kinetics of GLUT4 translocation leading to increased glucose uptake in response to key physiological stimulations should facilitate the discovery of modulators with therapeutic potentials.

## Results

### GLUT4 behavior in cultured human muscle cells

For these studies we used human SKM cells, a previously characterized large T antigen transformed myocyte cell line that can be differentiated *in vitro* into myoblasts and myotube-like cells^10^. To quantitatively assess GLUT4 behavior, we use a GLUT4 construct engineered with an HA-epitope in the first exofacial loop and GFP fused to its cytoplasmic carboxyl domain (HA-GLUT4-GFP), a reporter that has been extensively used in studies of GLUT4 traffic^17,18^. HA-GLUT4-GFP in the PM of individual cells is determined by anti-HA immunofluorescence (Cy3 fluorescence) of fixed intact cells, and the total amount of HA-GLUT4-GFP expressed per cell is determined by the GFP fluorescence. Ratiometric analyses of the Cy3-to-GFP fluorescence per cell is proportional to the fraction of total HA-GLUT4-GFP in the PM^17,19,20^. SKM cells were transduced with a lentivirus harboring HA-GLUT4-GFP cDNA to obtain a pooled population of cells stably expressing HA-GLUT4-GFP, hereafter referred to as SKM-GLUT4 cells. In vitro differentiated SKM-GLUT4 cells expressed myogenin and myosin heavy chain, two established muscle cell differentiation markers^21,22^. Incubation of cells in differentiation medium induced expression of these proteins within 3 days, and expression of these markers were maintained for at least 7 days post-differentiation (Figure 1A). The majority of SKM-GLUT4 cells formed multinucleated cells by day 3 of differentiation, although some cells remained mononuclear despite exposure to the differentiation protocol (Figure 1B). In both mononucleated and multinucleated cells, HA-GLUT4-GFP was distributed between the trans Golgi Network (**TGN**)/Golgi perinuclear region of cells and dispersed throughout the cytosol in puncta (vesicles). Unless noted otherwise, in the following discussions when we refer to the behavior of GLUT4 it is based on studies of HA-GLUT4-GFP. Because microscopy-based, single-cell methods were used to analyze the behavior of GLUT4, multinucleated and mononucleated cells were independently analyzed. The fraction of PM GLUT4 was reduced in differentiated cells compared to that in proliferative myocyte-like SKM-GLUT4 cells, with intracellular accumulation of HA-GLUT4-GFP being the greatest in multinucleated cells (Figure 1C). In multinucleated cells, 15.5 ± 2.0% of total GLUT4 (mean ± SEM) was on the PM. Intracellular retention of GLUT4 is a hallmark of differentiated muscle cells and adipocytes.

**Figure 1.**
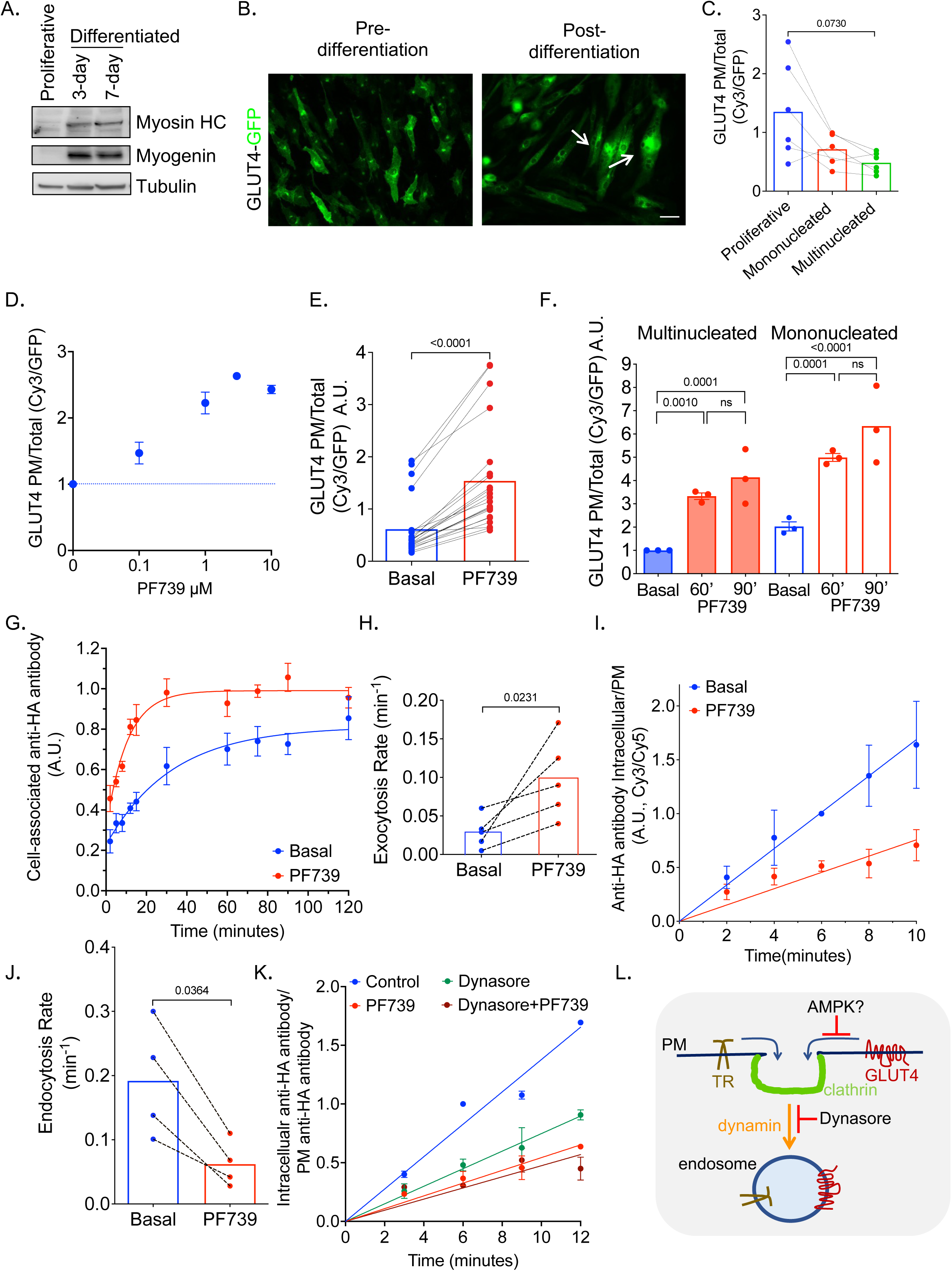
AMPK regulation of GLUT4 in differentiated human muscle cells. **A.** Western blot of markers of differentiation. **B.** Fluorescence microscopy imaging of GFP fluorescence of pre and post-differentiation (day 3) SKM cells stably expressing HA-GLUT4-GFP by lentiviral transduction. Arrows indicate multinucleated cells. Scale bar is 50µM. **C.** PM expression of HA-GLUT4-GFP. Each data point is the mean value calculated for of at least 40 cells per condition per experiment (n=6). The lines link the data for the 3 conditions from the individual experiments. p value, One-way ANOVA with unequal variances **D.** PF739 dose effect (60-minute treatment) on PM expression of HA-GLUT4-GFP in differentiated, multinucleated SKM cells. Each data point is the mean value calculated for of at least 40 cells per condition per experiment. Data are ± SD, n=2, normalized to the basal condition in each experiment. **E.** PM expression of HA-GLUT4-GFP in day 3 differentiated, multinucleated SKM-GLUT4 cells treated with 3 µM PF739 for 60 minutes. Each data point is the PF739 stimulated to unstimulated PM GLUT4 ratio calculated from the mean vales of at least 40 cells per condition in each of 23 independent experiments. p value, Mann-Whitney test. **F.** PM expression of HA-GLUT4-GFP in day-3 differentiated, multinucleated and mononucleated SKM-GLUT4 cells treated with or without 3 µM PF739 for 60 or 90 minutes. Basal condition is unstimulated cells. Each data point is the mean value calculated for at least 40 cells per condition per experiment. One-way ANOVA p<0.0001, p values are FDR multiple comparison adjustment. ns, non-significant. **G.** Exocytosis assay for HA-GLUT4-GFP in day-3 differentiated SKM-GLUT4 cells treated with or without 3 µM PF739 for 60 minutes. Data are cell-associated anti-HA as a function of incubation time in medium normalized to the average of 60, 75 and 90 min of basal (unstimulated) condition in each experiment. Data points are means ± SEM calculated for at least 40 cells per condition per experiment. n = 5. **H.** Exocytosis rate constants ± SEM determined from assays in panel G. Dashed lines join individual experiments. Bar graphs showing the mean. *p, S*tudent’s 2 sample t-test. **I.** Endocytosis assay for HA-GLUT4-GFP in day 3-differentiated SKM-GLUT4 cells treated with or without 3 µM PF739 for 60 minutes. Data are a ratio of internal anti-HA (internalized pulse of GLUT4) ratioed to PM GLUT4. Data are normalized to the 6-minute time point of basal (unstimulated) condition in each experiment. Mean values ± SEM, n = 4. **J.** Endocytosis rate constants ± SEM determined from assays in A. Dashed lines join individual experiments. Bar graphs showing the mean. *p, p, S*tudent’s 2 sample t-test. **K.** Endocytosis assay for HA-GLUT4-GFP in day 3-differentiated SKM-GLUT4 cells in basal (unstimulated) conditions, treated with 3 µM PF739 for 60 minutes, pre-treated with 100µM Dynasore for 60 min, or both PF739 and Dyansore for 60 min. PF739 and Dyansore were maintained in the media during the uptake measurements. Data are a ratio of internal GLUT4 to surface GLUT4, normalized to the 6-minute time point of basal condition in each experiment. Mean values ± SEM, n = 4. **L.** Endocytosis Cartoon. See text for description.

### AMPK activation promotes GLUT4 translocation to the PM of differentiated SKM cells

Treatment of SKM-GLUT4 cells with PF739, an AMPK activator non-selective for isoform composition^10^, stimulated phosphorylation of acetyl-Co carboxylase, a substrate of activated AMPK, to the same degree as 5-Aminoimidazole-4-carboxamide ribonucleoside (**AICAR**), a well-studied AMPK activator (Figure S1A). In a dose-dependent manner, a 60-minute PF739 stimulation of SKM-GLUT4 cells promoted an increase of GLUT4 in the PM (Figure 1D). A 2.9 ± 0.17-fold increase of PM GLUT4 in multinucleated SKM cells, relative to basal PM GLUT4, was achieved at 3 µM PF739 (Figure 1E). Both mono and multinucleated SKM cells responded to PF739 stimulation by translocation of GLUT4 to the PM (Figure 1F). Incubation of cells with PF739 for 90 minutes did not induce a greater increase in PM GLUT4, demonstrating a new steady state is reached within 60 minutes of AMPK stimulation (Figure 1F). All kinetic results reported hereafter are from analyses of multinucleated SKM cells stimulated with 3µM PF739 for 60 min.

### AMPK activation effects both GLUT4 exocytosis and endocytosis

To explore the mechanism underlying the change in PM GLUT4 following AMPK activation, we characterized GLUT4 exocytosis and endocytosis. To measure exocytosis, living cells are incubated with anti-HA in the medium and cell-associated anti-HA antibody is monitored as a function of time^19,20,23,24^. With increasing incubation times, cell-associated anti-HA antibody increases as unbound HA-GLUT4-GFP inside the cell traffics to the PM where it is bound by antibody. Cell-associated anti-HA antibody accumulation plateaus when all HA-GLUT4-GFP that is in equilibrium between intracellular compartments and the PM has been bound by the antibody (that is, has trafficked once to the PM). Thus, the plateau reflects the size of the GLUT4 pool that is in equilibrium with the PM, and the rate of rise to the plateau is how fast intracellular GLUT4 traffics to the PM (i.e., exocytosis rate constant).

In unstimulated SKM-GLUT4 cells, intracellular GLUT4 was constitutively exocytosed and endocytosed (Figure 1G-J). Thus, intracellular sequestration of GLUT4 in human SKM cells is a dynamic process determined by the kinetics of internalization and recycling. PF739 activation of AMPK stimulated an acceleration of GLUT4 exocytosis (Figure 1G). The GLUT4 exocytosis rate constant was increased by approximately three-fold in PF739-stimulated cells (Figure 1H). In basal SKM cells, cell-associated anti-HA plateaued at 87.7 ± 1.1% (mean ± SEM, n=5) of the plateau in PF739-stimulated cells, indicating that the effect of AMPK on GLUT4 exocytosis is predominantly to accelerate the movement of GLUT4 that is in equilibrium with the PM of unstimulated cells, although the small difference between the plateaus supports AMPK-dependent mobilization of a pool of GLUT4, constituting ∼12% of the cycling pool in stimulated cells, that is not in equilibrium with the PM in unstimulated cells.

To measure GLUT4 internalization, cells were pulse labelled with anti-HA antibody. A plot of the amount of internal anti-HA antibody ratioed to the amount of anti-HA bound to the PM yields a straight line whose slope is proportional to the internalization rate constant of HA-GLUT4-GFP^25,26^. PF739 activation of AMPK inhibited GLUT4 internalization by about a third (Figure 1I,J). Thus, AMPK controls PM amounts of GLUT4 by targeting both exocytosis and endocytosis. In rodent muscle cells, GLUT4 is internalized by dynamin-dependent mechanism^27^. We used Dynasore, a small molecule inhibitor of dynamin^28^, to investigate GLUT4 internalization in SKM cells. Dynasore inhibition of dynamin-mediated endocytosis in SKM-GLUT4 cells was confirmed by inhibition of transferrin receptor (**TR**) internalization, a client of dynamin-dependent endocytosis (Figure S1B). Acute inhibition of dynamin-dependent endocytosis by Dynasore inhibited GLUT4 endocytosis in SKM cells, consistent with a role for dynamin in GLUT4 internalization (Figure 1K). The effect of Dynasore on GLUT4 internalization was not additive to AMPK activation, providing evidence that AMPK targets dynamin-dependent internalization of GLUT4 (Figure 1K). However, AMPK inhibition of GLUT4 internalization is not due to a general inhibition of dynamin-dependent endocytosis because the internalization of TR was unaffected by PF739 activation of AMPK (Figure S1C). These data support a model in which AMPK regulates GLUT4 internalization at a step upstream of dynamin function in GLUT4 internalization, perhaps targeting GLUT4 clustering in clathrin-coated pits prior to endocytosis (Figure 1L).

### Insulin stimulation promotes GLUT4 translocation to the PM of differentiated SKM cells

Insulin control of GLUT4 trafficking has been extensively characterized in rodent muscle cell lines, where it has been shown to induce a near two-fold increase of PM GLUT4 by stimulating exocytosis^1^. Insulin (10 nM) stimulation of differentiated SKM-GLUT4 cells induced phosphorylation of the serine threonine kinase AKT and its downstream target TBC1D4 (AS160) (Figure S1D-F). TBC1D4 phosphorylation by AKT is required for GLUT4 translocation in adipocytes and muscle cells^24,29-31^. In multinucleated SKM cells, insulin stimulation induced a 51 ± 17% (mean ± SEM) increase of PM GLUT4 relative to that in unstimulated cells, whereas the effect of insulin on PM GLUT4 in mononucleated SKM cells was less (29 ± 14%) (Figure S1G). Insulin-stimulated redistribution of GLUT4 to the PM is accounted for by an acceleration in the exocytosis rate constant, 37 ± 11% (mean ± SD) increase, without a change in GLUT4 internalization (Figure S1H,I).

### TBC1D4-Rab10 signaling module link AMPK to regulation of GLUT4 exocytosis in SKM cells

Our results reveal fundamental differences between responses to insulin and AMPK in the control of GLUT4 in a model human muscle cells line, both in the magnitude of response and mechanisms underlying translocation. The Rab GAP TBC1D4 (AS160) is a key component of the machinery required for intracellular retention of GLUT4 in rodent adipocytes and muscle cells^24,29,32-34^. In humans, naturally occurring variants and mutations in TBC1D4 are associated with metabolic disruptions, reenforcing its role in regulating GLUT4 traffic to the PM^35,36^. CRISPR/CAS9 mediated knockout of TBC1D4 in SKM proliferative cells did not affect differentiation as assessed by formation of multinucleated cells (Figure S2A,B). There was a significant increase in PM GLUT4 in TBC1D4 knockout basal cells compared to control cells (Figure 2A). This increase was due to an approximate three-fold acceleration of GLUT4 exocytosis in unstimulated TBC1D4 KO SKM cells (Figure 2B,C). These data establish in human SKM cells that TBC1D4 functions to reduce PM GLUT4 in unstimulated conditions by inhibiting GLUT4 exocytosis, consistent with the results of previous studies in adipocytes and rodent muscle cell lines.

**Figure 2.**
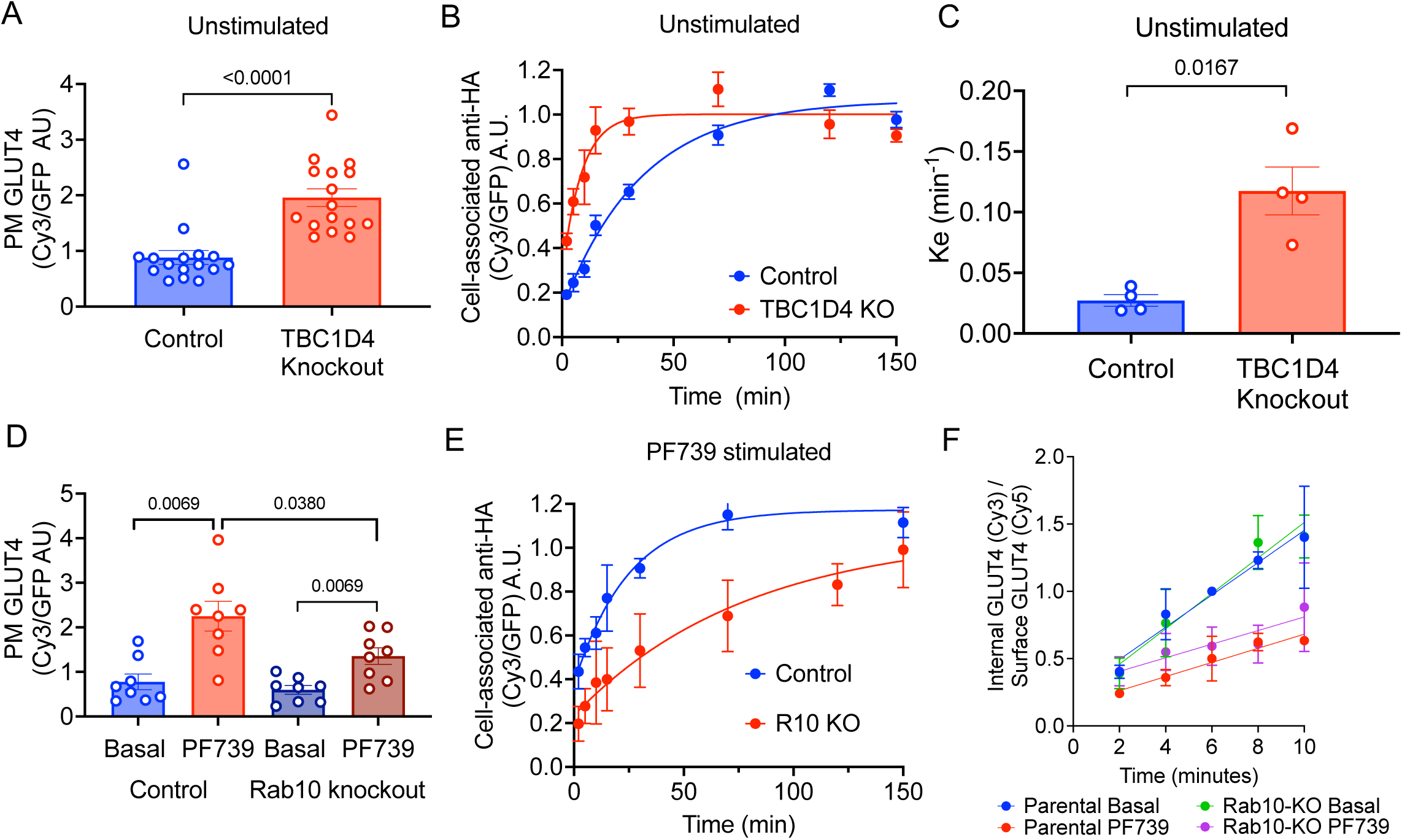
TBC1D4 and Rab10 are required for AMPK regulation of GLUT4 regulation of GLUT4. **A.** PM HA-GLUT4-GFP in day 3-differentiated SKM parental and TBC1D4 knockout (KO) cells stably expressing the HA-GLUT4-GFP, in basal (unstimulated) conditions. Each symbol is the mean value calculated from at least 40 cells in individual experiments (n=11). Data are presented as mean ± SEM. *p,* log Transformation Student’s 2 sample t-test **B.** Exocytosis assay for HA-GLUT4-GFP under basal (unstimulated) conditions in day 3-differentiated SKM-CRISPR/CAS9 parental and TBC1D4 KO cells stably expressing HA-GLUT4-GFP. Data are cell-associated anti-HA as a function of incubation time in medium containing anti-HA antibody, normalized to the average of the 70, 120 and 150 min of basal condition in each experiment. Data are mean values ± SEM, n = 6. **C.** Exocytosis rate constants ± SEM determined from assays in panel B. Mean ± SEM, n=4. *p,* Welch’s 2 sample t-test. **D.** PM HA-GLUT4-GFP in day 3-differentiated SKM parental cells and SKM-Rab10 knockout (KO) cells stably expressing HA-GLUT4-GFP, treated with 3 µM PF739 for 60 minutes. Each symbol is the mean value of at least 40 cells per experiment (n=9). Data are presented as mean ± SEM normalized to the basal condition of parental cells. p values, One-way ANOVA with unequal variances p=0.0009, FDR multiple comparison adjustment. **E.** Exocytosis assay for HA-GLUT4-GFP under basal (unstimulated) conditions in day 3-differentiated SKM parental and Rab10-KO cells stably expressing HA-GLUT4-GFP. Data are cell-associated anti-HA as a function of incubation time in medium containing anti-HA antibody, normalized to the average of 70, 120 and 150 min of basal condition in each experiment. Data are mean values ± SEM, n = 4. **F.** Endocytosis assay for HA-GLUT4-GFP in day 3-differentiated SKM-CRISPR/CAS9 parental cells and Rab10-KO SKM cells stably expressing GLUT4 treated with or without 3 µM PF739 for 60 minutes. Data are a ratio of internal anti-HA (internalized pulse of GLUT4) to PM GLUT4. Data are normalized to the 6-minute time point of basal (unstimulated) condition in each experiment. Mean values ± SEM, n = 3.

In unstimulated cells, TBC1D4 is an active GAP maintaining its target Rabs in the inactive GDP-bound form. TBC1D4 is phosphorylated by both AKT and AMPK^33,37^. Phosphorylation of TBC1D4 inactivates its GAP function, increasing the amount of its target Rabs in the GTP bound form. The targets of TBC1D4’s GAP activity are Rabs 2, 8a, 10 and 14^38^. Knockdown of Rab10 in cultured mouse adipocytes and Rab10 knockout in primary adipocytes (white and brown) reduces insulin stimulated GLUT4 translocation, identifying Rab10 as TBC1D4’s target required for GLUT4 translocation in adipocytes^13,14,39^. TBC1D4 is also an AMPK target; therefore, we investigated whether Rab10 has a role in AMPK induced translocation of GLUT4 in SKM cells. Knockout of Rab10 did not affect differentiation of SKM cells as assessed by formation of multinuclear cells (Figure S2C,D). PF739 stimulated translocation of GLUT4 to the PM was blunted in Rab10 knockout SKM cells (Figure 2D). The reduced translocation resulted from reduced PF739 stimulation of GLUT4 exocytosis in Rab10 knockout SKM cells (Figure 2E, Figure S2E). Rab10 knockout did not affect AMPK inhibition of GLUT4 internalization (Figure 2F).

It has been previously shown in rat muscle cells that Rab8a not Rab10 is the target of TBC1D4 required for insulin stimulated GLUT4 translocation^15,40^. Transient knockdown of Rab8a in differentiated SKM cells did not affect PF739 stimulated GLUT4 translocation to the PM (Figure S2F,G), whereas transient Rab10 knockdown blunted PF739 stimulated GLUT4 translocation, as expected based on the RAB10 knockout data (Figure S2H,I). Hence, as is the case for insulin stimulation of GLUT4 exocytosis in adipocytes, Rab10 is required for AMPK stimulation of GLUT4 exocytosis in human SKM cells. We were unable to interrogate the roles of Rab10 and Rab8a in insulin stimulated GLUT4 translocation in SKM cells because the smaller and more variable net effect of insulin on GLUT4 translocation, as compared to the effect of PF739.

### GLUT4 proximity proteome in unstimulated SKM cells

Having established that GLUT4 is dynamically retained intracellularly in unstimulated SKM cells, we next used APEX2 proximity mapping to further explore GLUT4’s trafficking itinerary in molecular detail. APEX2 is an engineered peroxidase whose substrate is biotin-phenol^41^. APEX2-generated biotin-phenol radical reacts with electron-dense amino acids to covalently derivatize proteins with biotin. Because the biotin-phenyl radical is highly unstable in aqueous solution, the reaction is limited to about 20nm from the site of its formation. Therefore, proteins within a radius of about 20nm of APEX2 tagged to GLUT4 will be biotinylated, and mass spectrometry of streptavidin-isolated biotinylated proteins can be used to identify those proteins. A GLUT4 construct in which the APEX2 peroxidase was fused to the cytosolic carboxyl terminus of GLUT4, replacing GFP of the HA-GLUT4-GFP construct (HA-GLUT4-APEX2), was stably expressed in SKM cells by lentiviral transduction (Figure S3A). HA-GLUT4-APEX2 stably expressed in SKM cells was localized to the perinuclear region of SKM cells and redistributed to the PM following PF739 stimulation (Figure S3B,C).

To catalog the GLUT4 proximal proteome, SKM cells were pre-incubated with APEX2 substrate biotin phenol for 60 minutes, followed by incubation with 0.38% H_2_O_2_ for 60 seconds. GLUT4 proximal proteins were identified by mass spectrometry analysis of proteins enriched by streptavidin-isolation. The GLUT4 proximity proteome was determined by comparing streptavidin-isolated proteins from cells incubated with biotin phenol and treated with H_2_O_2_ (complete reaction) to those isolated from cells treated with H_2_O_2_ without biotin phenol preincubation (no APEX2 substrate negative control). Analyses of the mass spectrometry intensities from 6 independent experiments yielded 700 proteins with log2(intensity full reaction/intensity no biotin phenol) > 0 and adjp < 0.05 (Benjamini-Hochberg false discovery corrected) (Table S1).

As anticipated, GLUT4 was identified as an APEX2 biotinylated protein enriched over the negative control in all six experiments (Figure 3A). The transmembrane insulin-regulated aminopeptidase (**IRAP**) was also identified as a component of the GLUT4 proximal proteome (Figure 3A). IRAP is the best described cargo of the insulin-regulated transport pathway that traffics GLUT4 to the PM in adipocytes and muscle cells^42^. In addition, endosomal cargo proteins: TR, low density lipoprotein (**LDL**) receptor and LDL receptor related protein 1 (**LRP1**) were identified as components of the GLUT4 proximal proteome (Figure 3A). All of these endosomal proteins have been described to localize, to varying degrees, with GLUT4 in adipocytes, despite these proteins not being primary cargos of the insulin-regulated pathway. The proximity of these proteins with GLUT4 detected by APEX2 mapping likely reflects the transit of GLUT4 through common endosomes during constitutive traffic to and from the PM in unstimulated SKM cells. Proteins previously shown to colocalize with GLUT4 in the adipocyte TGN Golgi retention compartment were significantly enriched in the GLUT4 basal proximal proteome, including those functionally required for GLUT4 traffic in adipocytes: sortilin^43-45^ cellugyrin^46^, GGA1^43,47^ and Sec16A^11,48^ as well as those with no known function in GLUT4 trafficking, ATP7A^11,49^ and trans Golgi network protein 2 (**TGN38**)^11,49^ (Figure 3A). Colocalization of GLUT4 with TGN38 and ATP7A were confirmed by immunofluorescence (Figure 3B). The overlap between GLUT4 proximal proteins in SKM muscle cells with those previously identified to colocalize with GLUT4 in adipocytes and rodent muscle cells support the hypothesis that GLUT4 traffic pathway is similar between muscle and adipocytes, and to a large degree, conserved between mouse and human cells.

**Figure 3.**
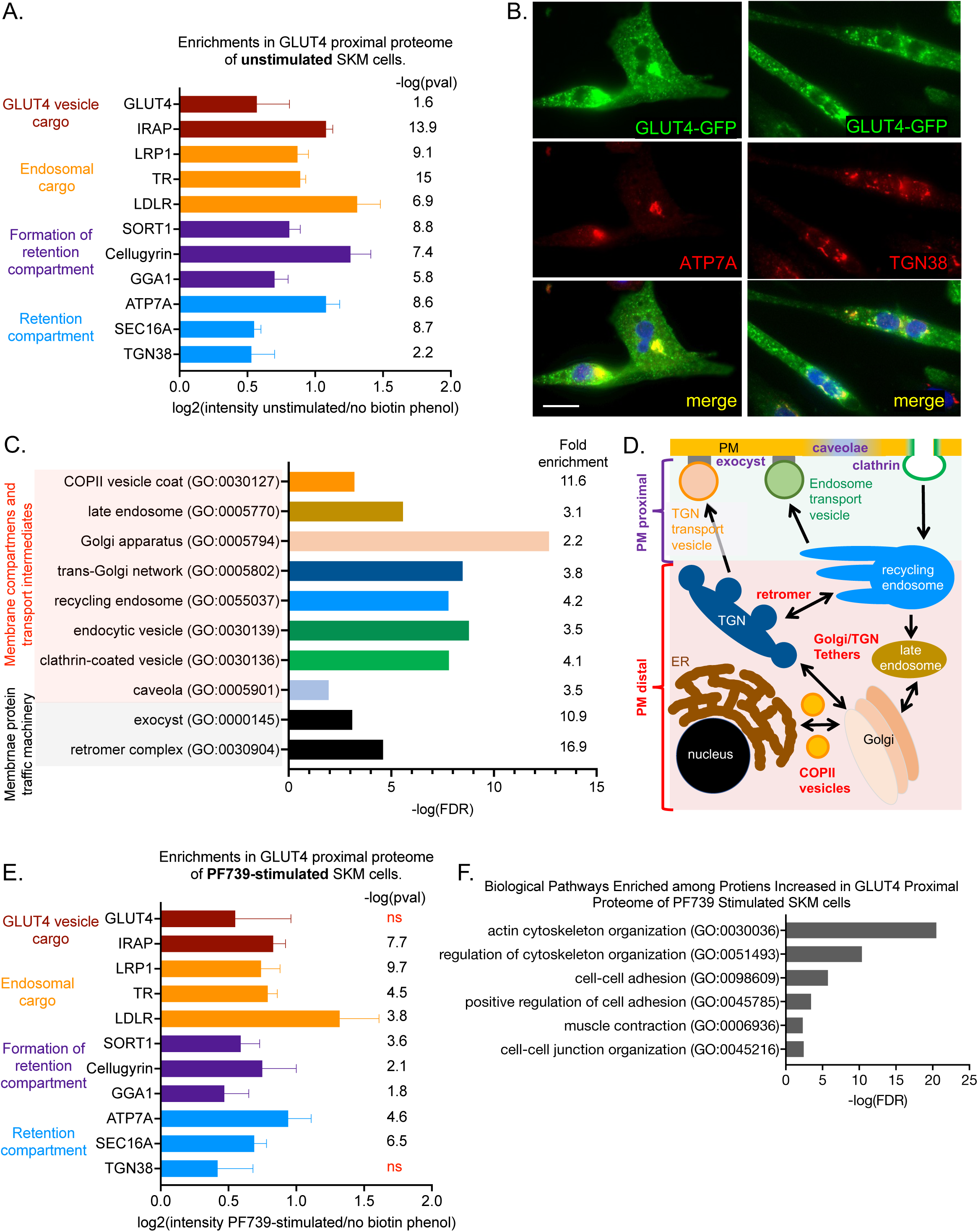
APEX2 proximal proteome mapping of GLUT4 in unstimulated and PF739 stimulated SKM differentiated cells. **A.** Enrichments of proteins in the unstimulated GLUT4 proximal proteome. Log2 values of the ratio of mass spectrometry intensities from experimental conditions to mass spectrometry intensities of reaction without biotin phenol, the APEX2 substrate. The p values are significant when tested against Benjamini-Hochberg correction for false discovery at alpha of 0.05. Data are from 6 experiments. **B.** Immunofluorescence co-colocalizations of ATP7A and TGN38 with HA-GLUT4-GFP in day 3 differentiated SKM cells. Scale bar is 50µM. **C.** Some biological groupings significantly overrepresented among proteins of GLUT4 proximal proteome in unstimulated cells. False discovery rate (FDR) calculated at alpha 0.05. Ontology pathway enrichment analyses were performed using Panther online software^50-52^. **D.** Schematic of compartments and protein complexes represented in the GLUT4 proximal proteome of SKM cells. See text for description. **E.** Enrichments of proteins in the PF739 stimulated GLUT4 proximal proteome. Log2 values of the ratio of mass spectrometry intensities from experimental conditions to mass spec intensities of reaction without biotin phenol, APEX2 substrate. The p values are significant when tested against Benjamini-Hochberg for false discovery at alpha of 0.05. ns, non-significant. Data are from 6 experiments. **F.** Some biological groupings significantly overrepresented among of proteins increased in the GLUT4 proximal proteome in PF739-stimulated cells compared to that in unstimulated cells. FDR calculated at alpha 0.05. Ontology pathway enrichment analyses were performed using Panther online software^50-52^.

Protein ontology pathway analyses^50-52^ established an enrichment of different intracellular compartments and transport pathways among the 700 proteins of the GLUT4 proximal proteome in unstimulated muscle cells (Figure 3C & D). These data reveal that in unstimulated SKM cells, GLUT4 was widely distributed among many intracellular compartments, providing independent support for our hypothesis that GLUT4’s intracellular retention is dynamically achieved in human muscle cells by constitutive transport of GLUT4 through various intracellular compartments, rather than by static sequestration within a single retention compartment. In addition, there were significant enrichments of cytoskeletal pathways, predominantly actin cytoskeleton, in the basal proteome (Figure S3D). Numerous previous studies have documented a role for the actin cytoskeleton in regulation of GLUT4 traffic in rodent muscle cells (reviewed in ref 1).

### GLUT4 translocation downstream of activated AMPK impacts the GLUT4 proximity proteome

We next explored how the GLUT4 proximal proteome is affected by AMPK activation. We focused on AMPK activation, rather than insulin stimulation given the robust GLUT4 translocation to the PM induced by PF739 as an AMPK activator and the desire to gain the mechanistic insight of the AMPK-mediated upregulation of glucose uptake that was previously less well understood. Of the proteins isolated by streptavidin precipitation, 399 had log2(intensity full reaction/no biotin phenol) > 0 and adjp < 0.05 (Benjamini-Hochberg false discovery corrected) for the comparison of 90 minutes PF739 stimulated (plus biotin phenol and H_2_O_2_) to no biotin phenol (negative control) (Table S2). All of these proteins were part of the GLUT4 proximal proteome of unstimulated cells. These data demonstrate that despite the markedly different kinetics of GLUT4 trafficking between unstimulated and PF739-stimulated cells, GLUT4 traverses many, if not all, of the same intracellular compartments in both states, thereby supporting the hypothesis that redistribution of GLUT4 to the PM is primarily achieved by altering traffic among these compartments rather than mobilizing a pool of GLUT4 that is static in unstimulated cells. Of note, IRAP, the endosomal proteins: LRP, TR and LDLR, and proteins associated with the perinuclear GLUT4 retention compartment: ATP7a, Sec16a, Sortilin, GGA and Cellugyrin, were all in the GLUT4 proximal proteome of AMPK activated cells (Figure 3E).

To quantify the effects of activated AMPK on relative abundances of proteins within the GLUT4 proteome, we contrasted the APEX2 mapping results of PF739-stimulated cells to APEX2 mapping generated in unstimulated cells. Of the 700 proteins of the GLUT4 proximity proteome in unstimulated cells, 526 were less abundant in the PF739-stimulated GLUT4 proximal proteome [log2(intensity PF739 stimulated cells/intensity unstimulated cells)] < 0, and 174 were more highly enriched [(log2(intensity PF739 stimulated cells/intensity unstimulated cells) > 0](Table S3). Proteins involved in the control of the actin cytoskeleton dynamics and functions, cell adhesion and cell-cell junction organization were significantly enriched among these 174 proteins (Figure 3F).

To further explore how the distribution of GLUT4 among the intracellular compartments was affected by PF739 activation of AMPK, we focused on protein complexes/groups known involved in membrane protein trafficking that were enriched in the GLUT4 proximal proteome of unstimulated cells (referenced to no biotin phenol control): exocyst proteins (Figure 4A*i*), caveolae proteins (Figure 4B*i*), clathrin triskelion proteins (Figure 4C*i*), Golgi/TGN peripheral proteins (Figure 4D*i*), COPII vesicle proteins (Figure 4E*i*) and retromer complex proteins (Figure 4F*i*). In each instance, multiple proteins of these complexes were significantly enriched, reinforcing the conclusion that GLUT4 is in proximity to these complexes in unstimulated cells. These complexes are also enriched in the GLUT4 proximal proteome generated in cells stimulated with PF739 (referenced to no biotin phenol control) (Figure 4A*ii,* B*ii,* C*ii,* D*ii,* E*ii* and *Fii*). The proximity of GLUT4 to these complexes in both stimulated and unstimulated cells, suggests the possibility that they participate in GLUT4 trafficking in both states.

**Figure 4.**
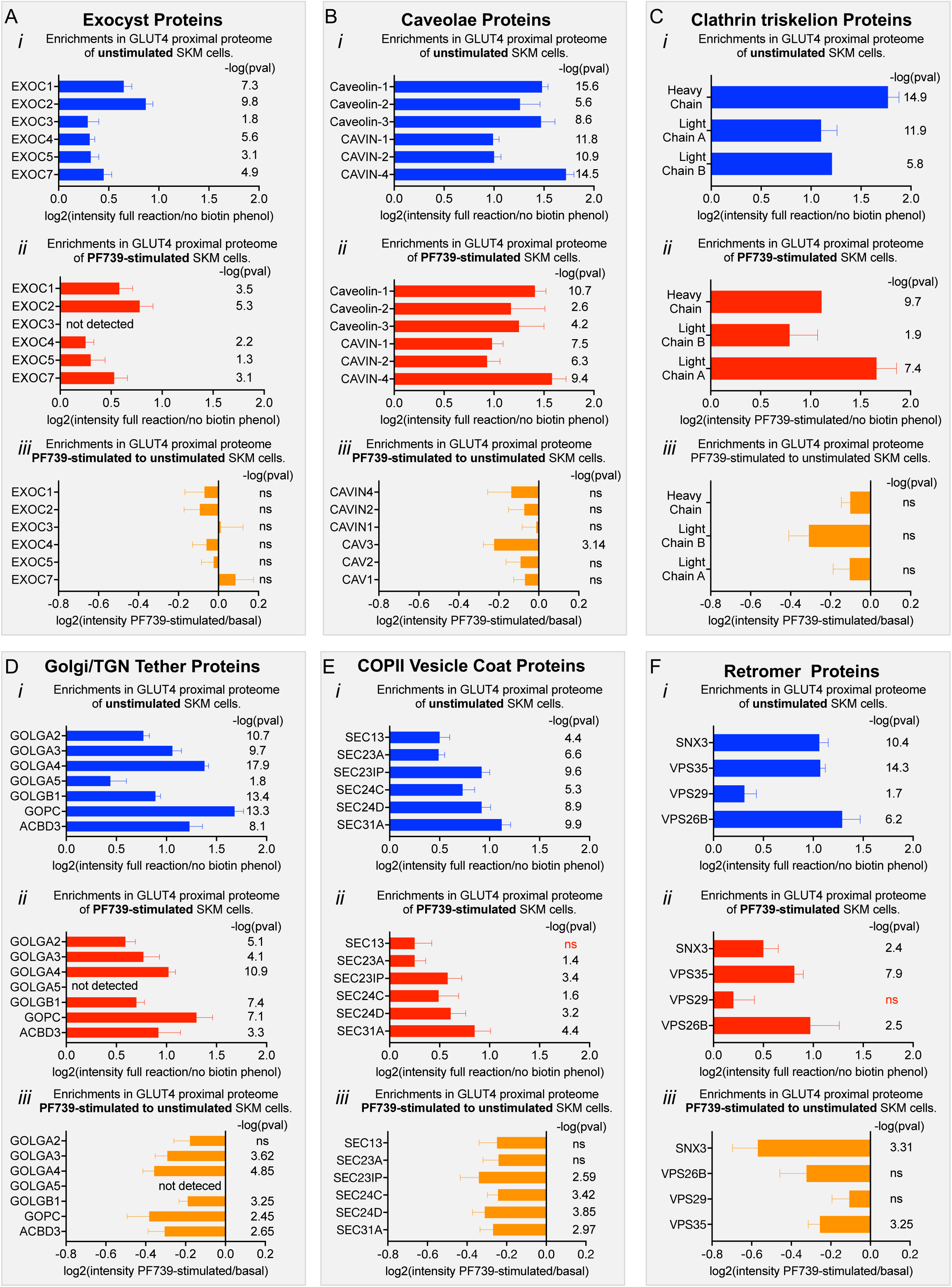
Impact of PF739 activation of AMPK on components of the GLUT4 proximal proteome. **A.** Exocyst proteins, **B.** Caveolae proteins, **C.** Clathrin triskelion proteins, **D.** Golgi/TGN tether proteins, **E.** COPII vesicle proteins, and **F.** Retromer proteins. Log2 of mass spectrometry ratios for i) unstimulated to no biotin phenol negative control, ii) PF739 stimulated to no biotin phenol negative control, and iii) PF739 stimulated to unstimulated calculated for each protein from 6 experiments are shown. ns, non-significant. The p values are significant when tested against Benjamini-Hochberg for false discovery at alpha of 0.05. ns, not significant at p< 0.05.

To explore whether the proximities of these proteins to GLUT4 were affected by AMPK activation, we contrasted the APEX2 biotinylation data from cells stimulated with PF739 to data from unstimulated cells. The mass spectrometry intensities of the exocyst (Figure 4A*iii*), caveolae (Figure 4B*iii*) and clathrin triskelion proteins (heavy and light chains) (Figure 4C*iii*) were not significantly different in PF739-stimulated compared to unstimulated SKM cells, indicating AMPK-activation and altered kinetics of GLUT4 trafficking had little difference on proximity of these protein complexes to GLUT4. The PF739-stimulated to unstimulated ratios of mass spectrometry intensities (log2) for Golgi/TGN tether proteins (Figure 4D*iii*), COPII vesicle proteins (Figure 4E*iii*) and retromer proteins (Figure 4F*iii*) were all negative, supporting the hypothesis that upon activation of AMPK, GLUT4 moves away from these proteins coincident with GLUT4 redistribution to the PM.

### Proximity proteome of GLUT4 trafficking mutant F^5^Y-GLUT4 is unaltered by activation AMPK activation

A GLUT4 mutant in which the phenylalanine of the amino terminal cytoplasmic domain FQQI motif (amino acids 5-8) has been mutated to tyrosine displays enhanced intracellular retention and a blunted insulin-stimulated translocation to the PM of adipocytes^11,26,53^. Despite the increased intracellular retention of F^5^Y-GLUT4 it is in equilibrium with the PM of unstimulated fat cells^11,26^. These data support a model in which the tyrosine substitution precludes insulin-stimulated mobilization of F^5^Y-GLUT4 to the PM by biasing F^5^Y-GLUT4 to the basal trafficking itinerary that is unaltered by insulin. We have previously used F^5^Y-GLUT4 as a tool to study GLUT4 trafficking in unstimulated adipocytes^11^.

F^5^Y-GLUT4-APEX2 stably (**F^5^Y-GLUT4**) expressed in SKM cells was better excluded from the PM than WT GLUT4, and AMPK-induced translocation of F^5^Y-GLUT4 to the PM was blocked (Figure S3D). Thus, the impact of the F^5^Y mutation on GLUT4 behavior in SKM muscle cells is similar to that in adipocytes: enhanced intracellular retention in unstimulated cells and blunted signal-induced translocation. We next used the F^5^Y-GLUT4-APEX2 to generate a proximal proteome in unstimulated and PF739 treated SKM cells. Analyses of the mass spectrometry intensities from 6 independent experiments yielded 836 proteins with unstimulated log2(intensity full reaction/no biotin phenol) > 0 and adjp < 0.05 (Benjamini-Hochberg false discovery corrected) for the comparison of complete reaction contrasted to reaction without biotin phenol as the negative control (Table S4). There was extensive overlap between the unstimulated proximal proteomes of F^5^Y-GLUT4 and WT GLUT4 (Figure 5A).

**Figure 5.**
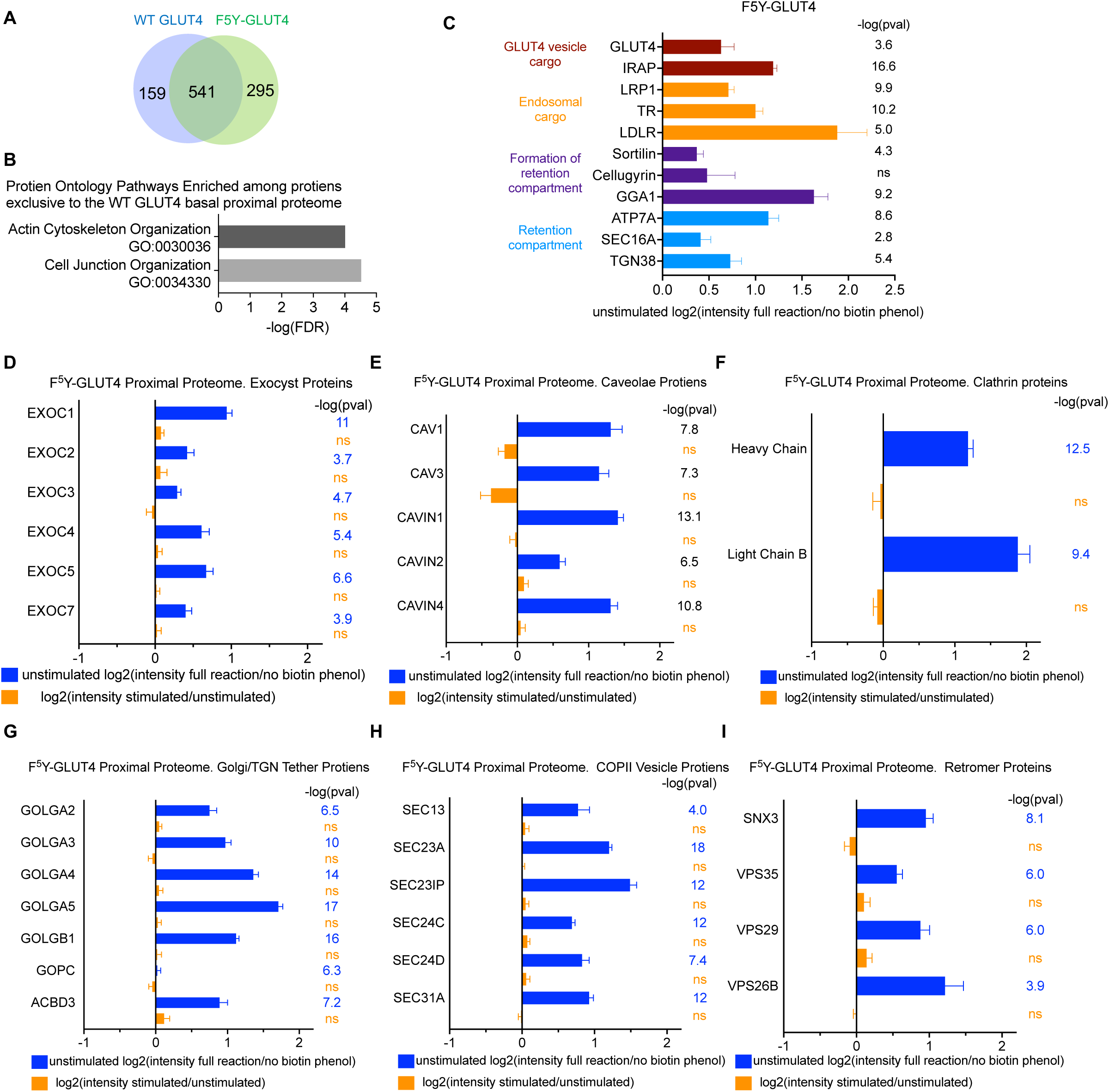
Impact of F^5^Ymutation in GLUT4, which affects GLUT4 trafficking, on GLUT4 proximal proteome. **A.** Venn diagram comparing the protein compositions of the unstimulated proximal proteomes of WT GLUT4 and F^5^Y-GLUT4. **B.** Some biological groupings significantly overrepresented among of proteins in the unstimulated WT GLUT4 proximal proteome that are not represented in the unstimulated F^5^Y-GLUT4 proximal proteome. FDR calculated at alpha 0.05. Ontology pathway enrichment analyses were performed using Panther online software^50-52^. **C.** Enrichments of proteins in the unstimulated F^5^Y-GLUT4 proximal proteome. Log2 values of the ratio of mass spectrometry intensities from experimental conditions to mass spec intensities of reaction without biotin phenol, APEX2 substrate. The p values shown are significant when tested against Benjamini-Hochberg for false discovery at alpha of 0.05. Data are from 6 experiments. **D.** Exocyst proteins, **E.** Caveolae proteins, **E.** Clathrin triskelion proteins, **G.** Golgi/TGN tether proteins, **H.** COPII vesicle proteins, and **I.** Retromer proteins. Log2 of mass spectrometry ratios for unstimulated to no biotin phenol negative control, (blue) and PF739-stimulated to unstimulated (orange) calculated for each protein from 6 experiments are shown. The p values shown are significant when tested against Benjamini-Hochberg for false discovery at alpha of 0.05.

Protein pathway ontology of the 295 proteins that were in the F^5^Y-GLUT4 proximal proteome but not the WT GLUT4 proximal proteome, did not reveal new compartments or reflect any biology specific to F^5^Y-GLUT4 (Figure S3E). These proteins were mostly within the same biology groupings as those identified for the WT GLUT4 proximal proteome, reinforcing the hypothesis that F^5^Y-GLUT4 is sequestered within the same compartments as WY GLUT4. Interestingly, there was an enrichment of actin cytoskeleton and cell junction proteins among the proteins in the WT but not F^5^Y-GLUT4 proximal proteome (Figure 5B). These findings are consistent the increased WT GLUT4 in the PM and corresponding reduced concentration in the TGN/Golgi compartments compared to F^5^Y-GLUT4, which would increase WT GLUT4’s proximity to actin and junctional proteins at or near the PM.

At the individual protein level, proteins enriched in GLUT4 vesicles and GLUT4-containing compartments in unstimulated SKM cells, were similarly enriched in the F^5^Y-GLUT4 proximal proteome (Figure 5C). In addition, the exocyst, caveolae, clathrin, Golgi/TGN tether proteins, COPII vesicle coat and retromer protein complexes and groups, which were enriched in the proximal proteome of WT GLUT4, were similarity enriched in the F^5^Y-GLUT4 proximal proteome of unstimulated SKM cells (Figure 5D-I). The enrichments of exocyst, caveolae and clathrin proteins in the F^5^Y-GLUT4 proximal proteome were not significantly reduced by AMPK stimulation, as was the case for WT GLUT4 (Figure 5D-F, Tables S5, S6). However, unlike the case for WT GLUT4, the enrichments of components of the perinuclear region of cells: Golgi/TGN tethers, COPII vesicle coat and retromer proteins in the F^5^Y-GLUT4 proximal proteome were not reduced by AMPK stimulation (Figure 5G-I). These data reinforce our hypothesis that translocation of GLUT4 to the PM primarily involves a redistribution from TGN/Golgi perinuclear region of cells, since the F^5^Y-GLUT4 is neither redistributed to the PM nor translocated away from the Golgi perinuclear region of the cells.

## Discussion

The phenomenon of GLUT4 translocation to the PM as a mechanism to regulate glucose homeostasis has been known for over 40 years. Yet despite important advances dissecting the molecular mechanisms regulating GLUT4 traffic to and from the PM of adipocytes and muscle cells, critical features of GLUT4 cell biology and its regulation by metabolic signaling pathways remain incompletely understood. One challenge in studying cell biology of GLUT4 is that its intracellular retention, translocation and subsequent sequestration following removal of the signal involves a number of cellular compartments, many of which are not specific to GLUT4 trafficking but are common to other transport processes. A detailed understanding of GLUT4’s itinerary and compartments traversed, both those common to other pathways and those more restricted to GLUT4, is required to understand molecular mechanisms that regulate the amount of GLUT4 in the PM, and consequently for a more complete description of the role of GLUT4 in whole-body metabolic regulation. APEX2 mapping provides a comprehensive view of a protein’s location. The GLUT4 proximal proteomes we have developed provide novel, comprehensive high resolution (∼20nM) views of the GLUT4 trafficking itinerary in human skeletal muscle cells and of how that itinerary is altered upon AMPK-induced GLUT4 translocation to the PM. This information on its own advances the field of GLUT4 cell biology by providing unbiased data in support for the dynamic retention model of GLUT4 retention and redistribution. In addition, the proximal proteome maps are a foundation for future studies of molecular mechanisms controlling GLUT4 traffic.

IRAP is known to follow the same trafficking pathway as GLUT4 in adipocytes and muscle cells and its enrichment in the GLUT4 proximal proteome provides validation for the data set. In addition, proteins previously shown to be required for generation of GLUT4 retention in unstimulated rodent adipocytes (Sortilin, GGA, and Cellugyrin), were enriched in the GLUT4 proximal proteome of unstimulated human muscle cells. Other proteins known to localize with or near GLUT4 in adipocytes -- ATP7a, Sec16A and TGN38 -- were also enriched in the proximal proteome of unstimulated human muscle cells. Thus, our novel findings reveal extensive overlap in GLUT4’s trafficking itinerary (defined by the proximal proteome atlas) from rodents to humans and between adipocytes and muscle cells.

From a mechanistic perspective, the diversity of endomembrane compartments represented in the GLUT4 proximal proteome of unstimulated human muscle cells supports a dynamic intracellular retention model in which GLUT4 continually recycles between the PM and intracellular compartments. The steady state accumulation in the PM reflects the overall rates of exocytosis and endocytosis, rather than static sequestration by tethering of GLUT4 within cells in the absence of signaling. The impact of AMPK activation on the GLUT4 proximal proteome further supports the redistribution among compartments accessible in unstimulated cells. Our dynamic retention and redistribution model is also supported by our kinetic studies in which we find constitutive endocytosis and exocytosis of GLUT4 in unstimulated and stimulated SKM cells. We have proposed a similar model for GLUT4 trafficking in adipocytes^11,19,23,48^. Our data supporting dynamic retention of GLUT4 in unstimulated human muscle cells agrees with results of studies of rodent muscle cells^1,2,54,55^. There are considerable precedents for steady state concentrations of membrane proteins within organelles/membrane compartments by active retrieval mechanisms, as we propose for GLUT4 (e.g.,^56^). However, an alternative model in which intracellular GLUT4 is not in equilibrium with the PM of unstimulated adipocytes and muscle cells and only traffics to the PM upon stimulation has been proposed^53,57-59^. Our kinetic data and APEX2 mapping data do not support a static retention model.

Importantly, we establish that AMPK activation induces GLUT4 translocation in human muscle cells by inhibition of endocytosis and acceleration of exocytosis. These data identify potential human and rodent differences because it has previously been shown that AMPK activation (either AICAR or ATP poison 2,4-dinitrophenol), induce GLUT4 translocation in L6 rat muscle cells by inhibition of endocytosis ^27,60^. In addition, we have also shown that the TBC1D4-Rab10 signaling module, which is key for insulin-induced translocation of GLUT4 to the PM of adipocytes, is required for AMPK-induced GLUT4 translocation in human muscle cells. Thus, the model we developed here can be used to further dissect the molecular mechanism linking physiologic signaling to the control of glucose uptake in human muscle cells.

Previous studies in adipocytes and rodent muscle cells have identified roles for the exocyst^61-63^, caveolae^64-66^, clathrin triskelion proteins^27,59^ and retromer^67,68^ in the control of GLUT4 trafficking in fat cells. Enrichments of these proteins in the GLUT4 proximal proteome of unstimulated cells highlight potential roles of the membrane transport steps controlled by these proteins in the steady state intracellular retention of GLUT4 in human muscle cells. The enrichments of COPII vesicle proteins and Golgi proteins suggest that in unstimulated muscle cells, GLUT4 cycles through or near both PM-proximal compartments and more PM-distal compartments (ER-Golgi/TGN region). These data, particularly the GLUT4 proximity to COPII vesicle coat proteins, agree with the hypothesized role for the ER Golgi Intermediate Compartment (**ERGIC**) in GLUT4 retention in muscle cells^69^. However, it is important to note that the proximity of GLUT4 with COPII vesicles does not mean GLUT4 is transported by COPII vesicles. Clathrin, in addition to having a role in GLUT4 internalization has also been reported to have a specific role in intracellular traffic of GLUT4 in human muscle controlled by a human-specific clathrin heavy chain isoform CHC22^70^; however, our analyses of the mass spectrometry data do not distinguish between CHC22 and the more conventional clathrin heavy chain CHC17.

Analyses of how AMPK activation alters the enrichments of these complexes in the GLUT4 proximal proteome revealed no significant change for the exocyst, caveolae and clathrin triskelion proteins, all of which are associated with more peripheral trafficking steps. Strikingly, the more PM distal aspects of cycling to and from the cell surface were reduced in AMPK stimulation, including retromer, Golgi tethers and COPII vesicle proteins. These data support a model in which AMPK-dependent acceleration of GLUT4 exocytosis is accompanied by a depletion of GLUT4 from the more distal compartments (that is, Golgi/TGN proximal) without a significant depletion from the PM proximal ones (Figure 4D). This link between reduced traffic of GLUT4 through Golgi proximal pathways and increased PM expression is also supported by the proximal proteome of F^5^Y-GLUT4. In this case, the blunted AMPK-induced PM translocation of F^5^Y-GLUT4 was associated with no redistribution of GLUT4 away from the Golgi proximal compartments. Reduced GLUT4 proximity to retromer, Golgi/TGN tether and COPII vesicle proteins in AMPK activated cells, are consistent with a more prominent role for these complexes in basal state retention. Nonetheless, they were also components of the GLUT4 proximal proteome of stimulated cells, establishing a similar GLUT4 itinerary in unstimulated and stimulated cells. Additional studies are required to functionally test the models of GLUT4 trafficking in human muscle cells that emerged from the APEX mapping data.

The most pronounced difference between the unstimulated and AMPK-stimulated GLUT4 proximal proteomes was an increase in proteins involved in the regulation of the actin cytoskeleton. These differences likely reflect a role for actin in the enhanced movement of GLUT4 to the PM of muscle cells, consistent with previous functional studies in rodent muscle cell lines^71-73^. The fewer proteins involved in actin biology in the F^5^Y-GLUT4 proximal proteome of unstimulated cells also highlights the role of the actin cytoskeleton in the constitutive transport of GLUT4 to the PM. The GLUT4 proximal proteome of PF739 stimulated cells did not support localization of GLUT4 to specific PM domains (or proximity to specific proteins) within the PM that might be linked to its function in increased glucose uptake, although such regulation might not be captured in the cultured cell model.

The TBC1D4-Rab10 signaling module is required for insulin-stimulated GLUT4 translocation in rodent adipocytes, with TBC1D4 promoting intracellular retention by inhibiting Rab10, and Rab10 promoting insulin-stimulated translocation^2^. GLUT4 exocytosis is the target of TBC1D4-Rab10. Here we show the TBC1D4-Rab10 signaling module is also required for regulation of GLUT4 traffic in human muscle cells: TBC1D4 for GLUT4 intracellular retention in basal adipocytes and Rab10 for GLUT4 translocation to the PM in response to AMPK. A major question in the field is to understand the impact of contraction/exercise and AMPK activation on insulin-stimulated GLUT4 translocation^74^.

Although the data presented here do not specifically address that question, the novel detailed information on GLUT4 itineraries in unstimulated and AMPK-stimulated cultured human muscle cells provide a foundation for future functional and mechanistic studies interrogating the intersection of signal transduction downstream of AMPK and insulin receptor.

## MATERIALS AND METHODS

### Chemicals, and drugs

Chemicals and drugs used were PF739 (Pfizer), Dynasore (S8047, Selleck Chemicals), Insulin (I5500, SigmaAldrich) and AICAR (M1404, Sigma-Aldrich).

### Cell culture and differentiation of human skeletal muscle cells

SKM cells^10^ were grown in muscle cell growth media (C-23060, PromoCell), supplemented with 2% horse serum (16050122, Thermofisher), 1% chick embryo extract (C3999, US Biologicals) and penicillin/streptomycin (15070063, Thermofisher) at 37°C/5% CO_2_. For differentiation, SKM cells were grown to confluency, the medium switched to differentiation medium containing Dulbecco’s Modified Eagle’s Medium low glucose, pyruvate (11885092, ThermoFisher), supplemented with 2% horse serum and penicillin/streptomycin. Unless noted otherwise, cells were studied at day 3 post differentiation.

### Generation of stable expression and CRISPR knockout cell lines

Engineered SKM cell lines were generated via lentiviral transduction. HEK293T packaging cells were transfected with the required lentiviral cDNA constructs using Lenti-X packaging system following the manufactures protocol (631276, Takara). SKM cells stably expressing HA-GLUT4-GFP (SKM-GLUT4 cells) were generated by transducing with HA-GLUT4-GFP in pLenti6 lentiviral vector. Following growth cells were sorted for GFP to isolate a pooled population of SKM cells stably expressing HA-GLUT4-GFP (SKM-GLUT4 cells). SKM-GLUT4 cells stably expressing CRISPR/CAS9 were generated by transducing SKM-GLUT4 with CRISPR/CAS9 lentivirus. Transduced cells were selected for stable CRISPR/CAS9 expression using hygromycin selection. Stable expression of CRISPR/CAS9 did not affect cell differentiation nor response to PF739. Rab10-KO and TBC1D4-KO cells were generated by transducing SKM-GLUT4-CRISPR/CAS9 cells with lentivirus containing gRNAs targeting Rab10 or TBC1D4 and selected for Zeomycin to enrich for cells expressing gRNAs. SKM-GLUT4-CRISPR/CAS9 cells were used as controls in studies of Rab10 and TBC1D4 knockout lines. gRNA sequencing targeting Rab10: CACCGCATTGCGCCTCTGTAGTAGG and targeting TBC1D4: CACCGGAAACAGGCCTTCAGTACGG were used.

### siRNA knockdowns in SKM cells

Day 3 differentiated SKM myotubes (one well of a six-well cluster) were trypsinized, pelleted, resuspended in 500 µl DMEM Pen/Strep and serum-free media, mixed with 2 nM of siRNA and electroporated at 180V and 950µF. Electroporated cells were plated in SKM differentiation media on coverslips, incubated for 3–4 hours in a 37°C/5% CO_2_ incubator, followed by addition of 2ml of SKM differentiation media. After 48–72 hours, cells were used for translocation assays, and for analyzing mRNA and protein expression. The siRNAs were purchased from Integrated DNA Technologies. The siRNA sequences are as follows: Rab10: si1: UAUGACAUCACCAAUGGUAAAAGTT; Rab8A: si1, GGCAAGAGAAUUAAACUGCAGAUAT;

### RNA isolation and quantitative real-time polymerase chain reaction (RT-PCR)

mRNA was isolated using the RNeasy kit (74106, QIAGEN) and cDNA was prepared using the RNA to cDNA EcoDry Premix (639545, Takara Bio). Quantitative RT-PCR was performed using appropriate primer pairs from the PrimerBank database.

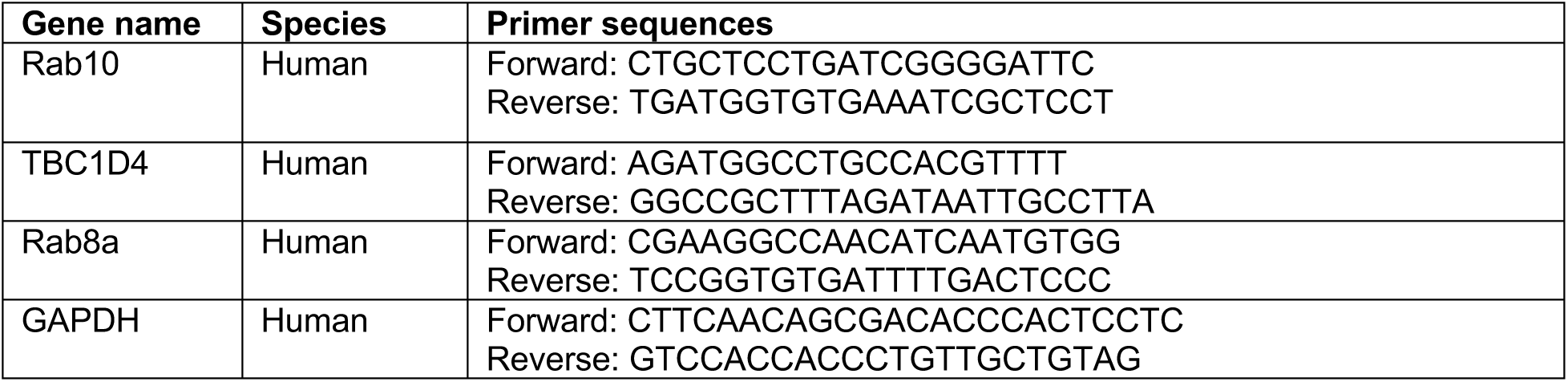

### Western blot analysis

Proteins were extracted from 3-day differentiated SKM cells using cell lysis buffer (9803S, Cell Signaling Technology), supplemented with a complete protease and phosphatase inhibitor cocktail (78442, ThermoFisher) and boiled at 95 °C for 5 minutes. Proteins displayed on were SDS-PAGE gels (8, 10 or 12% acrylamide, depending on the molecular weight of the protein to be visualized), transferred onto nitrocellulose membranes, and blocked at room temperature for 1 h in 5% milk or 5% BSA (for phospho-antibodies) in Tris-buffered saline-Tween (TBS-T). Membranes were then incubated with primary antibodies (diluted in 5% milk or 2.5% BSA with TBS-T) overnight at 4 °C. Primary antibodies used for Western blotting were against ß-tubulin (ab6046, Abcam), Rab10 (4262S, Cell Signaling Technology), Rab8A (610844, BD Biosciences), myosin heavy-chain (M4276, Millipore Sigma), myogenin (ab1835, Abcam), total acetyl-CoA carboxylase (ACC) (3676, Cell Signaling Technology), phospho-acetyl-CoA carboxylase (Ser79 ACC) (3661, Cell Signaling Technology), phospho T308-AKT (2965S, Cell Signaling Technology), total-AKT (9272S, Cell Signaling Technology), phospho-Thr642 TBC1D4 (8881S, Cell Signaling Technology), and total-TBC1D4 (07741, MilliporeSigma). Blots were incubated with goat anti-rabbit IgG-HRP or anti-mouse IgG-HRP secondary antibody (diluted 1:5000 in 5% milk or 2.5% BSA with TBS-T). Blots were visualized using My ECL Imager (Thermo Fisher) and analyzed using ImageStudioLite. Tubulin expression was used as a loading control where indicated.

### Microscopy

Images were acquired on an inverted epifluorescence microscope using a 20× air objective or a 40x oil objective (Leica Biosystems). Images were analyzed on MetaMorph (Universal Imaging) image processing software by appropriate thresholding and quantification, as previously described^17^. Signals for GFP and Cy3 were corrected for background and the surface (Cy3) to total (GFP) GLUT4 (S:T) was calculated per cell area, as previously detailed.

Airyscan images to acquire Z stack of the cells were collected using a laser scanning microscope (LSM880, ZEISS) with Airyscan using a 63X oil objective.

### GLUT4 translocation assay in SKM cells

HA-GLUT4-GFP translocation assay was performed as described previously^17^ with the following modifications. Day 3-differentiated SKM cells stably expressing HA-GLUT4-GFP were incubated in serum-free media for 2 h. Cells were stimulated with insulin or PF739 as specified in each experiment. Cells were then fixed in 3.7% formaldehyde for 6 min and PM HA-GLUT4-GFP detected by an anti-HA antibody (901503, BioLegend). HA staining was visualized by Cy3-goat anti-mouse secondary antibody (115-165-062, Jackson Immunoresearch). Cell nuclei were labeled with DAPI. Total HA-GLUT4-GFP was visualized by direct fluorescence of GFP. PM to total ratiometric values (dimensionless) were calculated by contrasting the PM HA signal intensity to the GFP signal per cell, after background correction for each channel^17^. The mean PM to total resulting values of 40 to 70 cells per condition were averaged and those averages across multiple independent experiments were compared. For graphical presentation of data across experiments, the values from individual experiments were normalized to a condition common to each experiment (for example, PM to total ratio of control unstimulated cells). Statistics were performed on non-normalized data.

### Exocytosis assay

Exocytosis assay was carried out as previously described^20^. Briefly, live cells were incubated in serum-free media at 37°C for 2 h, following which they were incubated in 2 ml of medium containing insulin or PF739 for 60 min to achieve new steady state of GLUT4 distribution. Cells were then incubated with a saturating concentration of anti-HA antibody (HA.11) (determined empirically for each batch of HA.11 antibody) serum-free DMEM supplemented with 1 mg/ml ovalbumin, for corresponding time points at 37°C/5% CO_2_, in the absence (control) or presence of the stimulant. Total cell-associated anti-HA antibody at specified times of incubation were determined in fixed, saponin-permeabilized cells by secondary staining with Cy3-goat anti-mouse antibody. Values of Cy3/GFP were calculated for multiple cells at individual time points and the mean values were plotted in Prism and fitted to a one-phase exponential equation. The slope of this curve provided the value of the rate of exocytosis (min^-1^). A separate set of dishes was used to quantify GLUT4 translocation PM-to-total ratio in the absence or presence of the particular stimulant, as a control to verify the effect of the stimulant. Cells were imaged and analyzed as described above.

### Endocytosis assay

GLUT4 endocytosis assay was carried out as previously described^20^. Briefly, live cells were preincubated in serum-free media at 37°C for 2 h, followed by incubation in 2 ml of media containing insulin or PF739 for 60 min to achieve new steady-state of GLUT4 distribution. Cells were then incubated with a saturating concentration of anti-HA antibody (HA.11) (determined empirically for each batch of HA.11 antibody) in serum-free DMEM supplemented with 1 mg/ml ovalbumin, for corresponding time points at 37°C/5% CO_2_, in the absence (control) or presence of the particular stimulant. Cells were washed, fixed and PM bound HA.11 stained with a saturating concentration of Cy5-goat anti-mouse antibody. Cells were then refixed, permeabilized with saponin, and stained with Cy3-goat anti-mouse antibody to reveal internal HA.11 (the Cy3 antibody binds to the same epitopes as the Cy5 antibody, and therefore does not bind to the surface-exposed epitopes already bound by the Cy5 antibody previously). The internal to PM value per cell was calculated for individual cells and averaged for each time point. A plot of that ratio versus time yields a straight-line whose slope is the internalization rate constant^20^.

TR internalization was assayed by incubating cells in serum-free DMEM containing 3µg/ml of Cy3-labelled-transferrin for specified times, in the absence (control) or presence of Dynasore (100µM) or PF739 (3 µM). Following fixation, cells were incubated with a saturating concentration of monoclonal anti-TR antibody (B3/25) in serum-free DMEM supplemented with 1 mg/ml ovalbumin, for corresponding time points at 37°C/5% CO_2_, Cells were then washed, fixed and surface-exposed TR was stained with a saturating concentration of Cy5-goat anti-mouse antibody, without permeabilizing the cells, to label the PM TR.

For both GLUT4 and TR internalization experiments, the internal to PM value per cell (Cy3/Cy5) was calculated for individual cells and averaged for each time point. A plot of that ratio versus time yields a straight-line whose slope is the internalization rate constant^20^. A separate set of dishes was used to quantify GLUT4 translocation surface-to-total ratio in the absence or presence of the particular stimulant, as a control to verify the effect of the stimulant.

### HA-GLUT4-APEX2

HA-GLUT4-APEX2 was created by replacing the GFP of HA-GLUT4-GFP in pLenti plasmid with APEX2 sequences and the resulting construct was confirmed by sequencing. HA-F^5^Y-GLUT4-APEX2 was created using Quikchange kit (Agilent, Inc) to change Phe^5^ to Tyr of HA-GLUT4-APEX2 in pLenti plasmid.

### HA-GLUT4-APEX2 translocation

Unlike measurement of PM HA-GLUT4-GFP, which is a single cell ratiometric determination using GFP fluorescence power to normalize the anti-HA signal per cell, measurement of PM HA-GLUT4-APEX2 is a cell population measurement in which the average PM anti-HA staining of cells (fixed, intact cells) from one dish is divided by the total HA-GLUT4-APEX2 expression measured in a separate dish by anti-HA immunofluorescence staining of fixed, permeabilized cells. This is necessary because the APEX2 construct does not have a second tag that can be used to normalize to total expression per cell. Because of needing to use population rather than single-cell normalized PM measures, the net translocation (that is, fold increase of PF739 stimulated over unstimulated) is smaller than that determined using ratiometric (that is, values corrected for expression per cell) measurements, thereby accounting for the apparently smaller translocation of HA-GLUT4-APEX2 than that of HA-GLUT4-GFP.

### APEX2 biotinylation

APEX2 proximity biotinylation assay was performed as previously described ^75^. Briefly, day 3-differentiated SKM cells stably expressing HA-GLUT4-APEX2 were incubated in serum-free media at 37°C for 2 hours followed by a 60 min incubation with or without biotin phenol (500 µM) in serum-free media. For PF739 stimulated conditions, PF739 was added to a final concentration of 3µM for the final 30 min of the two-hour incubation in serum-free media and 3 µM PF739 was maintained in the media during incubation with biotin phenol. Following the incubation with biotin-phenol, the cells were rapidly washed, placed on ice and incubated for 1 min with pre-chilled (4°C) 0.34% hydrogen peroxide (H_2_O_2_) in phosphate-buffered saline (PBS). Cells were washed three times with ice-cold quenching solution (5 mM Trolox, 10 mM Sodium Ascorbate, 10 mM Sodium Azide in 1x PBS), lysed in 1x RIPA lysis buffer (5 mM Trolox, 10 mM Sodium Ascorbate, 10 mM Sodium Azide, protease/phosphatase inhibitors in H_2_O) and protein concentration measured. Cell extracts were tumbled over night at 4°C with 200µl of Streptavidin agarose beads (Pierce) per 1mg of cell extract in 500µl final volume. Beads were washed twice with 1x RIPA lysis buffer, once with 1M KCl, once with 0.1 M NA_2_CO_3_, once with 2 M Urea in 10 mM Tris (pH 8.0), twice with 1x RIPA lysis buffer, twice with 50 mM Tris (pH 7.5), and twice with 2 M Urea in 50 mM Tris (pH 7.5).

### Proteomics analyses

#### Digestion and TMT labeling

The samples in 2 M Urea, 50 mM Tris pH 7.5 were treated with DL-dithiothreitol at a final concentration of 6.25mM for 30min at 25°C with shaking on a Thermomixer (ThermoFisher). Free cysteine residues were alkylated with 2-iodoacetamide at a final concentration of 34mM for 30min at 25°C while protected from light. LysC (1μg) was added, followed by incubation for 1h at 25°C with shaking on a Thermomixer. After incubation, the urea concentration was reduced to < 1M with the addition of 150μL of NH_4_HCO_3_. Finally, trypsin (2μg) was added, followed by incubation for 16h at 37°C with shaking on a Thermomixer.

After incubation, the digest was acidified with the addition of 5μL of formic acid, and samples were brought to 1mL with 0.1% formic acid. Peptides were desalted on C18 Sep-Pak cartridges (Waters, Milford, MA, USA). Briefly, cartridges were conditioned by sequential addition of i) Methanol, ii) 80% ACN/0.1% Formic Acid, iii) 0.1% Formic Acid, iv) 0.1% Formic Acid. All conditioning volumes were 800μL, and vacuum was used to elute the solutions. Following conditioning, the acidified peptide digest was loaded onto the cartridge with vacuum. The stationary phase was washed once with 800μL of 0.1% Formic Acid. Finally, samples were eluted from the cartridge using 300μL of 80% Acetonitrile/0.1% Formic Acid. Eluted peptides were dried under vacuum followed by reconstitution in water. Peptide yield was quantified by NanoDrop (ThermoFisher). An aliquot of 10μg of peptide per sample was transferred to a fresh Eppendorf tube, followed by chemical labeling with TMTPro 16plex reagent (ThermoFisher). The samples were divided in two TMTPro 16 plexes, with each plex containing three biological replicates per condition. All steps related to TMT labeling were carried out according to manufacturer’s instructions, including labeling, quenching, and combining of labeled samples. Once combined, a 20μg aliquot of combined/labeled peptide material was transferred to a fresh Eppendorf tube and dried under vacuum.

### Basic Reversed-Phase (High-pH) Fractionation and Sample Loading

The peptide aliquot was reconstituted in 300 μL of 0.1% Trifluoroacetic Acid, followed by fractionation on a High-pH/Reversed-Phase spin column (ThermoFisher) according to manufacturer’s instructions. The resulting 8 fractions were dried under vacuum. Each fraction was then reconstituted in 20μL of 5% Acetonitrile/0.1% Formic Acid, sonicated, and transferred to an autosampler vial. Vials were stored in the autosampler of an Easy-1200 nanoLC (ThermoFisher) maintained at 10°C.

### MS analyses

Peptides were separated on a 50 cm column composed of C18 stationary phase (Thermo Fisher ES903) using a gradient from 5% to 30% B over 2hr (Buffer A: 0.1% FA in HPLC grade water; Buffer B: 80% ACN, 0.1% FA) with a flow rate of 250nL/min using an EASY-nLC 1200 system (Thermo Fisher Scientific). MS data were acquired on a QE-HF mass spectrometer (Thermo Fisher Scientific) using a data-dependent acquisition top 10 method, AGC target 5e5, maximum injection time of 50 msec, scan range of 350-1800 m/z and a resolution of 60K. MS/MS was performed at a resolution of 60K, AGC target 1e5, maximum injection time of 105 msec, isolation window 0.7 m/z. Dynamic exclusion was set to 20 seconds.

All LC-MS/MS runs were analyzed using the Sequest algorithm (SEQUEST HT, Thermo Scientific) within Proteome Discoverer 2.4 (Thermo Scientific) against the UniProt human database. A 10 ppm MS1 error tolerance was used. Trypsin was set as the enzyme, allowing for 2 missed cleavages. TMT tags on lysine residues and peptide N-termini (229.162932 Da) and carbamidomethylation of cysteine residues (57.02146 Da) were set as static modifications, while oxidation of methionine residues (+15.99492 Da) and acetylation of peptide N-termini (42.011 Da) were set as variable modifications. Percolator was used as the false discovery rate calculator, and all peptides were filtered at the “Strict” Target FDR level of 0.01. Protein assembly was guided by principles of parsimony to produce the smallest set of proteins necessary to account for all observed peptides. Quantitation was performed in Proteome Discoverer with a reporter ion integration tolerance of 20 ppm for the most confident centroid within the “Reporter Ions Quantifier” node and the MS order was set to MS2. Reporter ion values were corrected for isotopic impurities using the manufacturer provided factors.

Proteomics workflow starts from the reporter ion intensities per feature reported in PD’s PSMs.txt file. When a peptide and charge combination is measured multiple times in a sample, only the maximum intensity is kept. The log2 peptide intensities are median normalized assuming equal input loading of all channels. Peptide intensities are summarized to protein intensities using Tukey’s median polish algorithm ^76^. Protein normalization using bridge channel was applied afterwards.

Ontology pathway enrichment analyses of the mass spectrometry data were performed using Panther online software^50-52^.

### Data Availability

The data that support the findings of this study are available from the corresponding author upon request. The mass spectrometry proteomics data have been deposited to the MassIVE repository (https://massive.ucsd.edu/) with the dataset identifier **MSV000092089**.

### Statistical analysis

#### Mass spectrometry

Outcomes are preprocessed log transformed proteins. A mixed effects model with appropriate fixed, random, correlation, and variance structure is used for all outcomes. If a random effect was not necessary, a generalized least squares linear model with appropriate fixed, correlation, and variance structure is used. All residuals were evaluated for meeting the normality assumption with the Shapiro-Wilks Test and Q-Q Plots and the homogeneity/sphericity assumption by residual plots. The log2 ratios of mass spectrometry intensities were tested for multiple testing false discovery using Benjamini-Hochberg approach. All analyses were performed with R version 3.5.1.

##### GLUT4 trafficking assays

Data analysis were performed using with R version 3.5.1 and graphed using Prism graph pad (v9.5.1). Results are expressed as mean ± SEM (standard error of mean) from at least 3 independent experiments, in which greater than 40 cells per condition were analyzed. Statistical analyses are specified in the legends. All statistically analyses were done using raw, non-normalized data, although in some instances, for presentation purposes, normalized data are shown. Values of *p <*0.05 were considered to be statistically significant.

## Author contributions

AR: Performed experiments, analyzed data and wrote draft of manuscript.

JW: Performed experiments and analyzed data.

LY: Performed experiments, analyzed data, and edited manuscript.

JC: Performed mass spectrometry.

JG: mass spectrometry data pipeline

LX: mass spectrometry data pipeline

KH: Reviewed data and edited manuscript

BBZ: Reviewed data and edited manuscript

MJB: Reviewed data and edited manuscript

QX: Reviewed data, generated cell lines and edited manuscript

MM: Designed experiments, analyzed data and edited manuscript.

TEM: Conceived of project, designed experiments, analyzed data and edited manuscript.

## Acknowledgements

We thank members of the McGraw lab for discussion and critical reading of the manuscript and Amira Klip for helpful comments and suggestions.

## Figure Legends

**Supplemental Figure S1 in support of Figure 1.**
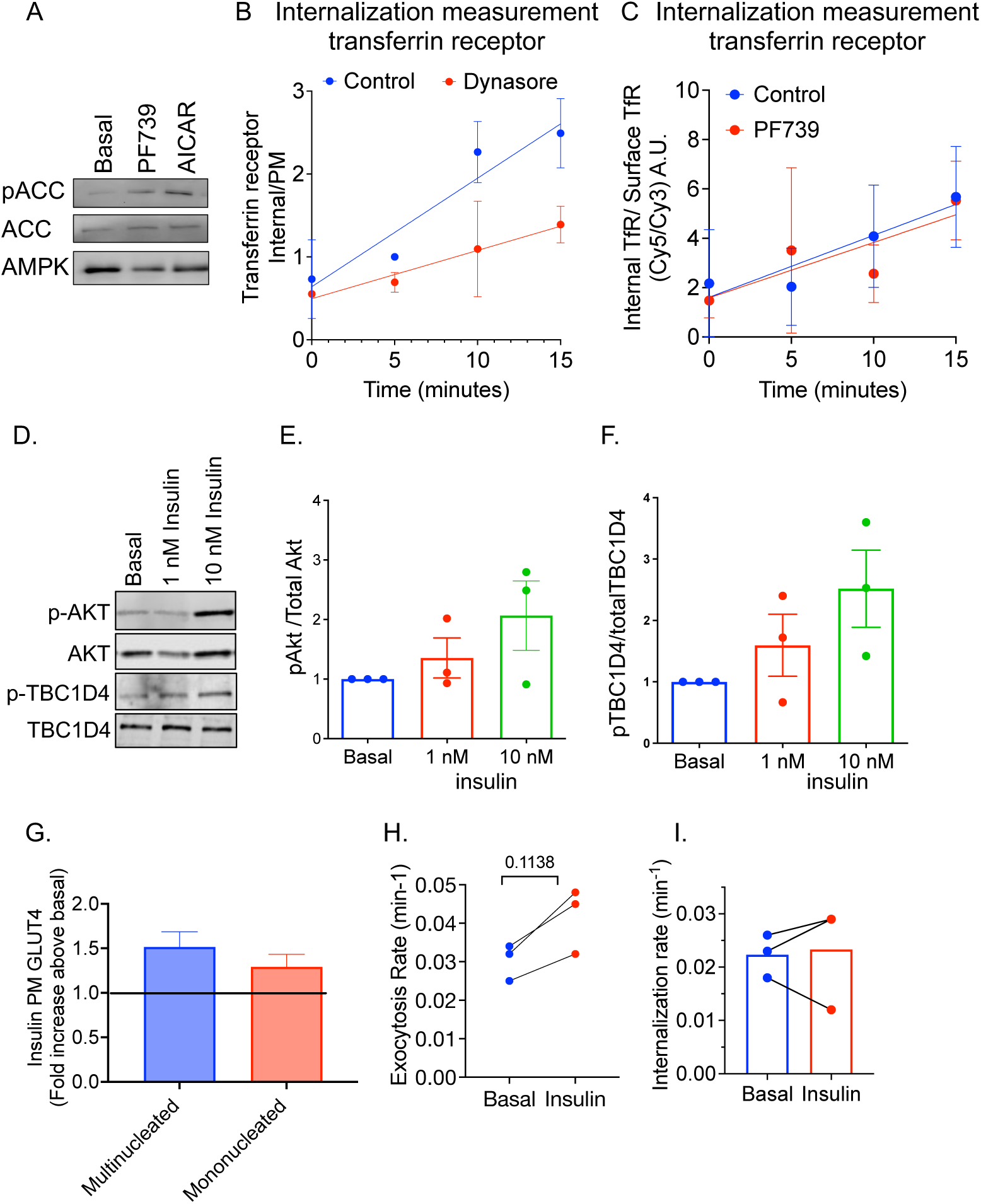
**A.** Western blot of day-3 differentiated SKM-GLUT4 cells treated for 60 min with vehicle, 3µM PF739 or 1 mM AICAR for phospho ACC (Ser79), total ACC, and AMPK. **B.** Effect of 100µM Dynasore on transferrin receptor (TR) internalization. Internalization rate constant is the slope of the line. Symbols are the mean ± SD of data from 3 independent experiments. Data are normalized to the 5-min data point of the control experiment of the individual experiments. **C.** Effect of 3 µM PF739 on TR internalization. Internalization rate constant is the slope of the line. Data are from a representative experiment. Each symbol is the mean value of at least 40 cells ± SD. **D.** Western blot analysis of pAkt (The308) and pAS160 (T642) in day 3-differentiated SKM-GLUT4 cells under basal and insulin stimulated (1 nM and 10 nM for 30 minutes) conditions. **E., F.** Quantification of phosphoblots from 3 independent experiments like that shown in panel D. ***G.*** Quantification of PM HA-GLUT4-GFP in day-3 differentiated SKM-GLUT4 cells (mononucleated and multinucleated). Serum starved cells were treated without (basal) or with 10 nM insulin for 30 minutes. The mean fold increase in PM HA-GLUT4-GFP above unstimulated cells determined from 40-70 cells were per condition were calculated in 16-18 independent experiments. Data are presented as mean ± SEM of the independent experiments. log Transformation Student’s 2 sample t-test p=0.0159, FDR multiple comparison adjustment (mono.basal vs mono.Insulin, p= 0.4418; multi.basal vs multi.Insulin.10 p= 0.2695). ***H.*** Exocytosis rate constants determined from HA-GLUT4-GFP exocytosis assays in day-3 differentiated SKM-GLUT4 cells in basal (unstimulated) and with 10 nM insulin stimulation for 30 minutes. Dashed lines join individual experiments. Bar graphs showing the mean. *p* value, Student’s 2 sample t-test. **I.** Endocytosis rate constants determined from HA-GLUT4-GFP endocytosis assays in day-3 differentiated SKM-GLUT4 cells, in basal (unstimulated) and with 10 nM insulin stimulation for 30 minutes. Dashed lines join data of individual experiments. Bar graphs show the mean. ns, non-significant using Student’s paired two-tailed t-test.

**Supplemental Figure S2 in support of Figure 2.**
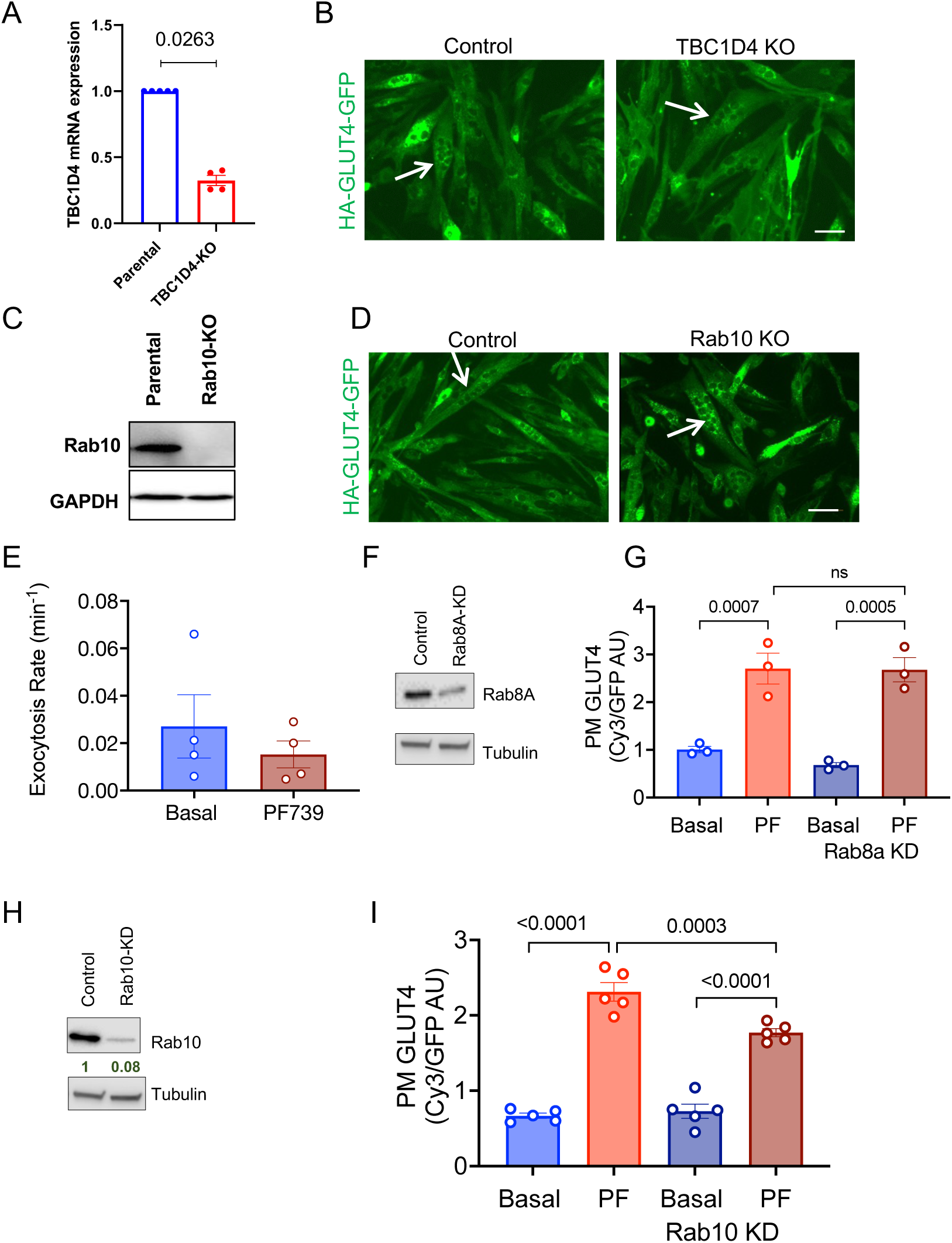
**A.** TBC1D4 mRNA expression in TBC1D4 KO cells. Two-sided paired Student’s T test on non-normalized data. Student’s 2 sample t-test. **B.** Fluorescence imaging for HA-GLUT4-GFP in differentiated SKM-CRISPR/CAS9 (control) and TBC1D4 KO SKM cells. Arrows note multinuclear cells. Scale bar 20µm. **C.** Assessment of Rab10 knockout by western blotting. **D.** Fluorescence imaging for HA-GLUT4-GFP in differentiated SKM-CRISPR/CAS9 (control) and Rab10 KO SKM cells. Arrows indicate multinuclear cells. Scale bar 20µm. **E**. GLUT4 rate constants measured in 4 independent experiments of control and Rab10 KO cells in unstimulated and PF739-stimulated conditions. **F.** Rab8a knockdown (KD) measured by western blotting. **G.** PM GLUT4 in unstimulated and PF739 stimulated control and Rab8a KD cells. Each symbol are data from independent experiments. One-way ANOVA p= 0.0002, p values FDR multiple comparison adjustment. **H.** Rab10 KD in SKM cells measured by western blotting. **I.** PM GLUT4 in unstimulated and PF739 stimulated control and Rab10 KD cells. Each symbol are data from independent experiments. PM GLUT4 in unstimulated and PF739 stimulated control and Rab8a KD cells. Each symbol are data from independent experiments. One-way ANOVA p < 0.0001, p values FDR multiple comparison adjustment.

**Supplemental Figure S3 in support of Figure 3.**
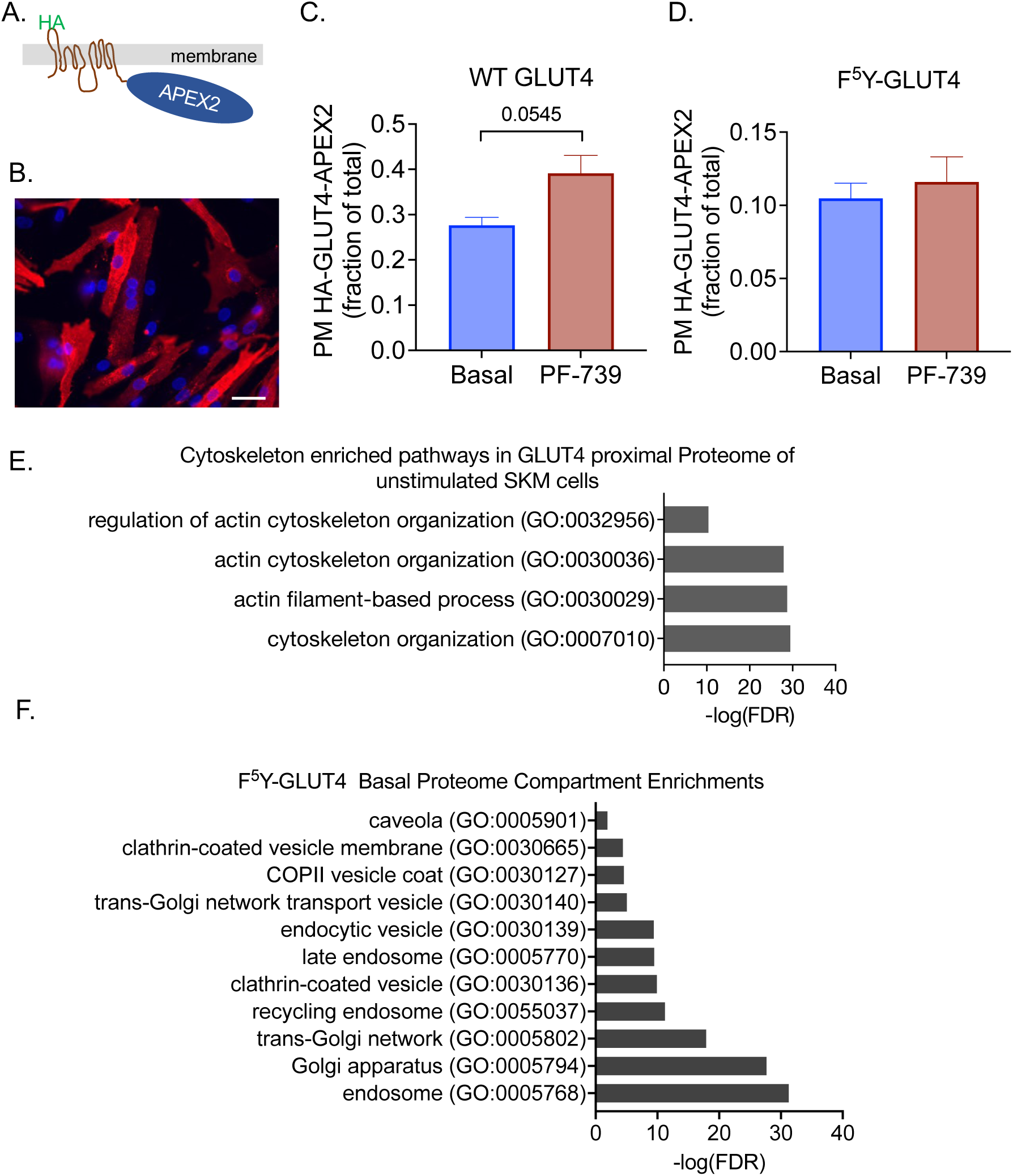
**A.** Cartoon of HA-GLUT4-APEX2. APEX2 replaces GFP of HA-GLUT4-GFP. **B.** Immunofluorescence of permeabilized SKM cells stably expressing HA-GLUT4-APEX2 stained with anti-HA antibody. **C.** Translocation of HA-GLUT4-APEX2 to the PM of SKM cells by PF739 activation of AMPK. Unlike measurement of PM HA-GLUT4-GFP, which is a single cell ratiometric determination using GFP fluorescence power to normalize the anti-HA signal per cell, measurement of PM HA-GLUT4-APEX2 is a cell population measurement in which the average PM anti-HA staining of cells (fixed, intact cells) from one dish is divided by the total HA-GLUT4-APEX2 expression measured in a separate dish by anti-HA IF staining of fixed, permeabilized cells. This is necessary because the APEX2 construct does not have a second tag that can be used to normalize to total expression per cell. Because these are a population rather than single cell normalized PM measures, the net translocation (that is, fold increase of PF739-stimulated over unstimulated) is smaller than that determined using ratiometric measurements (that is, values corrected for expression per cell), thereby accounting for the apparently smaller translocation of HA-GLUT4-APEX2 than HA-GLUT4-GFP. Welch’s 2-sample t-test. **D.** Translocation of HA-F^5^Y-GLUT4-APEX2 to PM of SKM cells measured as discussed for panel C. **E.** Some cytoskeleton pathways highly enriched among proteins of the GLUT4 proximal proteome in unstimulated cells. **F.** Some pathways highly enriched among proteins of the F^5^Y-GLUT4 proximal proteome in unstimulated cells. Ontology enrichment analyses were performed using Panther online software^79-81^.

## References

1. Sylow, L., Tokarz, V.L., Richter, E.A., and Klip, A. (2021). The many actions of insulin in skeletal muscle, the paramount tissue determining glycemia. Cell Metab 33, 758–780. 10.1016/j.cmet.2021.03.020.

2. Klip, A., McGraw, T.E., and James, D.E. (2019). Thirty sweet years of GLUT4. J Biol Chem 294, 11369–11381. 10.1074/jbc.REV119.008351.

3. Santoro, A., McGraw, T.E., and Kahn, B.B. (2021). Insulin action in adipocytes, adipose remodeling, and systemic effects. Cell Metab 33, 748–757. 10.1016/j.cmet.2021.03.019.

4. Richter, E.A., and Hargreaves, M. (2013). Exercise, GLUT4, and skeletal muscle glucose uptake. Physiol Rev 93, 993-1017. 10.1152/physrev.00038.2012.

5. Bird, S.R., and Hawley, J.A. (2016). Update on the effects of physical activity on insulin sensitivity in humans. BMJ Open Sport Exerc Med 2, e000143. 10.1136/bmjsem-2016-000143.

6. Jeon, S.M. (2016). Regulation and function of AMPK in physiology and diseases. Exp Mol Med 48, e245. 10.1038/emm.2016.81.

7. Kjøbsted, R., Hingst, J.R., Fentz, J., Foretz, M., Sanz, M.N., Pehmøller, C., Shum, M., Marette, A., Mounier, R., Treebak, J.T., et al. (2018). AMPK in skeletal muscle function and metabolism. FASEB J 32, 1741–1777. 10.1096/fj.201700442R.

8. Sylow, L., Kleinert, M., Richter, E.A., and Jensen, T.E. (2017). Exercise-stimulated glucose uptake - regulation and implications for glycaemic control. Nat Rev Endocrinol 13, 133–148. 10.1038/nrendo.2016.162.

9. Kjobsted, R., Munk-Hansen, N., Birk, J.B., Foretz, M., Viollet, B., Bjornholm, M., Zierath, J.R., Treebak, J.T., and Wojtaszewski, J.F. (2017). Enhanced Muscle Insulin Sensitivity After Contraction/Exercise Is Mediated by AMPK. Diabetes 66, 598–612. 10.2337/db16-0530.

10. Cokorinos, E.C., Delmore, J., Reyes, A.R., Albuquerque, B., Kjøbsted, R., Jørgensen, N.O., Tran, J.L., Jatkar, A., Cialdea, K., Esquejo, R.M., et al. (2017). Activation of Skeletal Muscle AMPK Promotes Glucose Disposal and Glucose Lowering in Non-human Primates and Mice. Cell Metab 25, 1147–1159.e1110. 10.1016/j.cmet.2017.04.010.

11. Brumfield, A., Chaudhary, N., Molle, D., Wen, J., Graumann, J., and McGraw, T.E. (2021). Insulin-promoted mobilization of GLUT4 from a perinuclear storage site requires RAB10. Mol Biol Cell 32, 57–73. 10.1091/mbc.E20-06-0356.

12. Sadacca, L.A., Bruno, J., Wen, J., Xiong, W., and McGraw, T.E. (2013). Specialized sorting of GLUT4 and its recruitment to the cell surface are independently regulated by distinct Rabs. Mol Biol Cell 24, 2544–2557. 10.1091/mbc.E13-02-0103.

13. Sano, H., Eguez, L., Teruel, M.N., Fukuda, M., Chuang, T.D., Chavez, J.A., Lienhard, G.E., and McGraw, T.E. (2007). Rab10, a target of the AS160 Rab GAP, is required for insulin-stimulated translocation of GLUT4 to the adipocyte plasma membrane. Cell Metab 5, 293–303. 10.1016/j.cmet.2007.03.001.

14. Vazirani, R.P., Verma, A., Sadacca, L.A., Buckman, M.S., Picatoste, B., Beg, M., Torsitano, C., Bruno, J.H., Patel, R.T., Simonyte, K., et al. (2016). Disruption of Adipose Rab10-Dependent Insulin Signaling Causes Hepatic Insulin Resistance. Diabetes 65, 1577–1589. 10.2337/db15-1128.

15. Sun, Y., Bilan, P.J., Liu, Z., and Klip, A. (2010). Rab8A and Rab13 are activated by insulin and regulate GLUT4 translocation in muscle cells. Proc Natl Acad Sci U S A 107, 19909–19914. 10.1073/pnas.1009523107.

16. Li, Z., Yue, Y., Hu, F., Zhang, C., Ma, X., Li, N., Qiu, L., Fu, M., Chen, L., Yao, Z., et al. (2018). Electrical pulse stimulation induces GLUT4 translocation in C(2)C(12) myotubes that depends on Rab8A, Rab13, and Rab14. Am J Physiol Endocrinol Metab 314, E478-E493. 10.1152/ajpendo.00103.2017.

17. Lampson, M.A., Schmoranzer, J., Zeigerer, A., Simon, S.M., and McGraw, T.E. (2001). Insulin-regulated release from the endosomal recycling compartment is regulated by budding of specialized vesicles. Mol Biol Cell 12, 3489–3501. 10.1091/mbc.12.11.3489.

18. Dawson, K., Aviles-Hernandez, A., Cushman, S.W., and Malide, D. (2001). Insulin-regulated trafficking of dual-labeled glucose transporter 4 in primary rat adipose cells. Biochem Biophys Res Commun 287, 445–454. 10.1006/bbrc.2001.5620.

19. Karylowski, O., Zeigerer, A., Cohen, A., and McGraw, T.E. (2004). GLUT4 is retained by an intracellular cycle of vesicle formation and fusion with endosomes. Mol Biol Cell 15, 870–882. 10.1091/mbc.e03-07-0517.

20. Blot, V., and McGraw, T.E. (2008). Use of quantitative immunofluorescence microscopy to study intracellular trafficking: studies of the GLUT4 glucose transporter. Methods Mol Biol 457, 347–366. 10.1007/978-1-59745-261-8_26.

21. Venuti, J.M., Morris, J.H., Vivian, J.L., Olson, E.N., and Klein, W.H. (1995). Myogenin is required for late but not early aspects of myogenesis during mouse development. J Cell Biol 128, 563–576. 10.1083/jcb.128.4.563.

22. Miano, J.M., Cserjesi, P., Ligon, K.L., Periasamy, M., and Olson, E.N. (1994). Smooth muscle myosin heavy chain exclusively marks the smooth muscle lineage during mouse embryogenesis. Circ Res 75, 803–812. 10.1161/01.res.75.5.803.

23. Martin, O.J., Lee, A., and McGraw, T.E. (2006). GLUT4 distribution between the plasma membrane and the intracellular compartments is maintained by an insulin-modulated bipartite dynamic mechanism. J Biol Chem 281, 484–490. 10.1074/jbc.M505944200.

24. Eguez, L., Lee, A., Chavez, J.A., Miinea, C.P., Kane, S., Lienhard, G.E., and McGraw, T.E. (2005). Full intracellular retention of GLUT4 requires AS160 Rab GTPase activating protein. Cell Metab 2, 263–272. 10.1016/j.cmet.2005.09.005.

25. Blot, V., and McGraw, T.E. (2006). GLUT4 is internalized by a cholesterol-dependent nystatin-sensitive mechanism inhibited by insulin. EMBO J 25, 5648–5658. 10.1038/sj.emboj.7601462.

26. Blot, V., and McGraw, T.E. (2008). Molecular mechanisms controlling GLUT4 intracellular retention. Mol Biol Cell 19, 3477–3487. 10.1091/mbc.e08-03-0236.

27. Antonescu, C.N., Diaz, M., Femia, G., Planas, J.V., and Klip, A. (2008). Clathrin-dependent and independent endocytosis of glucose transporter 4 (GLUT4) in myoblasts: regulation by mitochondrial uncoupling. Traffic 9, 1173–1190. 10.1111/j.1600-0854.2008.00755.x.

28. Kirchhausen, T., Macia, E., and Pelish, H.E. (2008). Use of dynasore, the small molecule inhibitor of dynamin, in the regulation of endocytosis. Methods Enzymol 438, 77–93. 10.1016/S0076-6879(07)38006-3.

29. Zeigerer, A., McBrayer, M.K., and McGraw, T.E. (2004). Insulin stimulation of GLUT4 exocytosis, but not its inhibition of endocytosis, is dependent on RabGAP AS160. Mol Biol Cell 15, 4406–4415. 10.1091/mbc.e04-04-0333.

30. Thong, F.S., Bilan, P.J., and Klip, A. (2007). The Rab GTPase-activating protein AS160 integrates Akt, protein kinase C, and AMP-activated protein kinase signals regulating GLUT4 traffic. Diabetes 56, 414–423. 10.2337/db06-0900.

31. Cartee, G.D. (2015). Roles of TBC1D1 and TBC1D4 in insulin- and exercise-stimulated glucose transport of skeletal muscle. Diabetologia 58, 19–30. 10.1007/s00125-014-3395-5.

32. Dokas, J., Chadt, A., Nolden, T., Himmelbauer, H., Zierath, J.R., Joost, H.G., and Al-Hasani, H. (2013). Conventional knockout of Tbc1d1 in mice impairs insulin- and AICAR-stimulated glucose uptake in skeletal muscle. Endocrinology 154, 3502–3514. 10.1210/en.2012-2147.

33. Espelage, L., Al-Hasani, H., and Chadt, A. (2020). RabGAPs in skeletal muscle function and exercise. J Mol Endocrinol 64, R1–R19. 10.1530/JME-19-0143.

34. Lansey, M.N., Walker, N.N., Hargett, S.R., Stevens, J.R., and Keller, S.R. (2012). Deletion of Rab GAP AS160 modifies glucose uptake and GLUT4 translocation in primary skeletal muscles and adipocytes and impairs glucose homeostasis. Am J Physiol Endocrinol Metab 303, E1273–1286. 10.1152/ajpendo.00316.2012.

35. Moltke, I., Grarup, N., Jorgensen, M.E., Bjerregaard, P., Treebak, J.T., Fumagalli, M., Korneliussen, T.S., Andersen, M.A., Nielsen, T.S., Krarup, N.T., et al. (2014). A common Greenlandic TBC1D4 variant confers muscle insulin resistance and type 2 diabetes. Nature 512, 190–193. 10.1038/nature13425.

36. Dash, S., Sano, H., Rochford, J.J., Semple, R.K., Yeo, G., Hyden, C.S., Soos, M.A., Clark, J., Rodin, A., Langenberg, C., et al. (2009). A truncation mutation in TBC1D4 in a family with acanthosis nigricans and postprandial hyperinsulinemia. Proc Natl Acad Sci U S A 106, 9350–9355. 10.1073/pnas.0900909106.

37. Treebak, J.T., Pehmøller, C., Kristensen, J.M., Kjøbsted, R., Birk, J.B., Schjerling, P., Richter, E.A., Goodyear, L.J., and Wojtaszewski, J.F. (2014). Acute exercise and physiological insulin induce distinct phosphorylation signatures on TBC1D1 and TBC1D4 proteins in human skeletal muscle. J Physiol 592, 351–375. 10.1113/jphysiol.2013.266338.

38. Miinea, C.P., Sano, H., Kane, S., Sano, E., Fukuda, M., Peranen, J., Lane, W.S., and Lienhard, G.E. (2005). AS160, the Akt substrate regulating GLUT4 translocation, has a functional Rab GTPase-activating protein domain. Biochem J 391, 87–93. 10.1042/BJ20050887.

39. Picatoste, B., Yammine, L., Leahey, R.A., Soares, D., Johnson, E.F., Cohen, P., and McGraw, T.E. (2021). Defective insulin-stimulated GLUT4 translocation in brown adipocytes induces systemic glucose homeostasis dysregulation independent of thermogenesis in female mice. Mol Metab 53, 101305. 10.1016/j.molmet.2021.101305.

40. Ishikura, S., and Klip, A. (2008). Muscle cells engage Rab8A and myosin Vb in insulin-dependent GLUT4 translocation. Am J Physiol Cell Physiol 295, C1016–1025. 10.1152/ajpcell.00277.2008.

41. Lam, S.S., Martell, J.D., Kamer, K.J., Deerinck, T.J., Ellisman, M.H., Mootha, V.K., and Ting, A.Y. (2015). Directed evolution of APEX2 for electron microscopy and proximity labeling. Nat Methods 12, 51–54. 10.1038/nmeth.3179.

42. Descamps, D., Evnouchidou, I., Caillens, V., Drajac, C., Riffault, S., van Endert, P., and Saveanu, L. (2020). The Role of Insulin Regulated Aminopeptidase in Endocytic Trafficking and Receptor Signaling in Immune Cells. Front Mol Biosci 7, 583556. 10.3389/fmolb.2020.583556.

43. Shi, J., Huang, G., and Kandror, K.V. (2008). Self-assembly of Glut4 storage vesicles during differentiation of 3T3-L1 adipocytes. J Biol Chem 283, 30311–30321. 10.1074/jbc.M805182200.

44. Shi, J., and Kandror, K.V. (2005). Sortilin is essential and sufficient for the formation of Glut4 storage vesicles in 3T3-L1 adipocytes. Dev Cell 9, 99–108. 10.1016/j.devcel.2005.04.004.

45. Shi, J., and Kandror, K.V. (2007). The luminal Vps10p domain of sortilin plays the predominant role in targeting to insulin-responsive Glut4-containing vesicles. J Biol Chem 282, 9008–9016. 10.1074/jbc.M608971200.

46. Kupriyanova, T.A., and Kandror, K.V. (2000). Cellugyrin is a marker for a distinct population of intracellular Glut4-containing vesicles. J Biol Chem 275, 36263–36268. 10.1074/jbc.M002797200.

47. Watson, R.T., Khan, A.H., Furukawa, M., Hou, J.C., Li, L., Kanzaki, M., Okada, S., Kandror, K.V., and Pessin, J.E. (2004). Entry of newly synthesized GLUT4 into the insulin-responsive storage compartment is GGA dependent. EMBO J 23, 2059–2070. 10.1038/sj.emboj.7600159.

48. Bruno, J., Brumfield, A., Chaudhary, N., Iaea, D., and McGraw, T.E. (2016). SEC16A is a RAB10 effector required for insulin-stimulated GLUT4 trafficking in adipocytes. J Cell Biol 214, 61–76. 10.1083/jcb.201509052.

49. Fazakerley, D.J., Naghiloo, S., Chaudhuri, R., Koumanov, F., Burchfield, J.G., Thomas, K.C., Krycer, J.R., Prior, M.J., Parker, B.L., Murrow, B.A., et al. (2015). Proteomic Analysis of GLUT4 Storage Vesicles Reveals Tumor Suppressor Candidate 5 (TUSC5) as a Novel Regulator of Insulin Action in Adipocytes. J Biol Chem 290, 23528–23542. 10.1074/jbc.M115.657361.

50. Ashburner, M., Ball, C.A., Blake, J.A., Botstein, D., Butler, H., Cherry, J.M., Davis, A.P., Dolinski, K., Dwight, S.S., Eppig, J.T., et al. (2000). Gene ontology: tool for the unification of biology. The Gene Ontology Consortium. Nat Genet 25, 25–29. 10.1038/75556.

51. Mi, H., Muruganujan, A., Ebert, D., Huang, X., and Thomas, P.D. (2019). PANTHER version 14: more genomes, a new PANTHER GO-slim and improvements in enrichment analysis tools. Nucleic Acids Res 47, D419–D426. 10.1093/nar/gky1038.

52. Mi, H., Muruganujan, A., Huang, X., Ebert, D., Mills, C., Guo, X., and Thomas, P.D. (2019). Protocol Update for large-scale genome and gene function analysis with the PANTHER classification system (v.14.0). Nat Protoc 14, 703-721. 10.1038/s41596-019-0128-8.

53. Govers, R., Coster, A.C., and James, D.E. (2004). Insulin increases cell surface GLUT4 levels by dose dependently discharging GLUT4 into a cell surface recycling pathway. Mol Cell Biol 24, 6456–6466. 10.1128/MCB.24.14.6456-6466.2004.

54. Randhawa, V.K., Thong, F.S., Lim, D.Y., Li, D., Garg, R.R., Rudge, R., Galli, T., Rudich, A., and Klip, A. (2004). Insulin and hypertonicity recruit GLUT4 to the plasma membrane of muscle cells by using N-ethylmaleimide-sensitive factor-dependent SNARE mechanisms but different v-SNAREs: role of TI-VAMP. Mol Biol Cell 15, 5565–5573. 10.1091/mbc.e04-03-0266.

55. Li, D., Randhawa, V.K., Patel, N., Hayashi, M., and Klip, A. (2001). Hyperosmolarity reduces GLUT4 endocytosis and increases its exocytosis from a VAMP2-independent pool in l6 muscle cells. J Biol Chem 276, 22883–22891. 10.1074/jbc.M010143200.

56. Banfield, D.K. (2011). Mechanisms of protein retention in the Golgi. Cold Spring Harb Perspect Biol 3, a005264. 10.1101/cshperspect.a005264.

57. Coster, A.C., Govers, R., and James, D.E. (2004). Insulin stimulates the entry of GLUT4 into the endosomal recycling pathway by a quantal mechanism. Traffic 5, 763–771. 10.1111/j.1600-0854.2004.00218.x.

58. Meriin, A.B., Zaarur, N., Bogan, J.S., and Kandror, K.V. (2022). Inhibitors of RNA and protein synthesis cause Glut4 translocation and increase glucose uptake in adipocytes. Sci Rep 12, 15640. 10.1038/s41598-022-19534-5.

59. Bogan, J.S., and Kandror, K.V. (2010). Biogenesis and regulation of insulin-responsive vesicles containing GLUT4. Curr Opin Cell Biol 22, 506–512. 10.1016/j.ceb.2010.03.012.

60. Fazakerley, D.J., Holman, G.D., Marley, A., James, D.E., Stockli, J., and Coster, A.C. (2010). Kinetic evidence for unique regulation of GLUT4 trafficking by insulin and AMP-activated protein kinase activators in L6 myotubes. J Biol Chem 285, 1653–1660. 10.1074/jbc.M109.051185.

61. Wang, S., Crisman, L., Miller, J., Datta, I., Gulbranson, D.R., Tian, Y., Yin, Q., Yu, H., and Shen, J. (2019). Inducible Exoc7/Exo70 knockout reveals a critical role of the exocyst in insulin-regulated GLUT4 exocytosis. J Biol Chem 294, 19988–19996. 10.1074/jbc.RA119.010821.

62. Inoue, M., Chang, L., Hwang, J., Chiang, S.H., and Saltiel, A.R. (2003). The exocyst complex is required for targeting of Glut4 to the plasma membrane by insulin. Nature 422, 629–633. 10.1038/nature01533.

63. Fujimoto, B.A., Young, M., Carter, L., Pang, A.P.S., Corley, M.J., Fogelgren, B., and Polgar, N. (2019). The exocyst complex regulates insulin-stimulated glucose uptake of skeletal muscle cells. Am J Physiol Endocrinol Metab 317, E957–E972. 10.1152/ajpendo.00109.2019.

64. Kandror, K.V., Stephens, J.M., and Pilch, P.F. (1995). Expression and compartmentalization of caveolin in adipose cells: coordinate regulation with and structural segregation from GLUT4. J Cell Biol 129, 999–1006. 10.1083/jcb.129.4.999.

65. Fecchi, K., Volonte, D., Hezel, M.P., Schmeck, K., and Galbiati, F. (2006). Spatial and temporal regulation of GLUT4 translocation by flotillin-1 and caveolin-3 in skeletal muscle cells. FASEB J 20, 705–707. 10.1096/fj.05-4661fje.

66. Gonzalez-Munoz, E., Lopez-Iglesias, C., Calvo, M., Palacin, M., Zorzano, A., and Camps, M. (2009). Caveolin-1 loss of function accelerates glucose transporter 4 and insulin receptor degradation in 3T3-L1 adipocytes. Endocrinology 150, 3493–3502. 10.1210/en.2008-1520.

67. Pan, X., Zaarur, N., Singh, M., Morin, P., and Kandror, K.V. (2017). Sortilin and retromer mediate retrograde transport of Glut4 in 3T3-L1 adipocytes. Mol Biol Cell 28, 1667–1675. 10.1091/mbc.E16-11-0777.

68. Yang, Z., Hong, L.K., Follett, J., Wabitsch, M., Hamilton, N.A., Collins, B.M., Bugarcic, A., and Teasdale, R.D. (2016). Functional characterization of retromer in GLUT4 storage vesicle formation and adipocyte differentiation. FASEB J 30, 1037–1050. 10.1096/fj.15-274704.

69. Camus, S.M., Camus, M.D., Figueras-Novoa, C., Boncompain, G., Sadacca, L.A., Esk, C., Bigot, A., Gould, G.W., Kioumourtzoglou, D., Perez, F., et al. (2020). CHC22 clathrin mediates traffic from early secretory compartments for human GLUT4 pathway biogenesis. J Cell Biol 219. 10.1083/jcb.201812135.

70. Gould, G.W., Brodsky, F.M., and Bryant, N.J. (2020). Building GLUT4 Vesicles: CHC22 Clathrin’s Human Touch. Trends Cell Biol 30, 705–719. 10.1016/j.tcb.2020.05.007.

71. Moller, L.L.V., Klip, A., and Sylow, L. (2019). Rho GTPases-Emerging Regulators of Glucose Homeostasis and Metabolic Health. Cells 8. 10.3390/cells8050434.

72. Tong, P., Khayat, Z.A., Huang, C., Patel, N., Ueyama, A., and Klip, A. (2001). Insulin-induced cortical actin remodeling promotes GLUT4 insertion at muscle cell membrane ruffles. J Clin Invest 108, 371–381. 10.1172/JCI12348.

73. Talior-Volodarsky, I., Randhawa, V.K., Zaid, H., and Klip, A. (2008). Alpha-actinin-4 is selectively required for insulin-induced GLUT4 translocation. J Biol Chem 283, 25115–25123. 10.1074/jbc.M801750200.

74. Jensen, J., and O’Rahilly, S. (2017). AMPK is required for exercise to enhance insulin sensitivity in skeletal muscles. Mol Metab 6, 315–316. 10.1016/j.molmet.2017.01.012.

75. Hung, V., Udeshi, N.D., Lam, S.S., Loh, K.H., Cox, K.J., Pedram, K., Carr, S.A., and Ting, A.Y. (2016). Spatially resolved proteomic mapping in living cells with the engineered peroxidase APEX2. Nat Protoc 11, 456–475. 10.1038/nprot.2016.018.

76. Huang, T., Choi, M., Tzouros, M., Golling, S., Pandya, N.J., Banfai, B., Dunkley, T., and Vitek, O. (2020). MSstatsTMT: Statistical Detection of Differentially Abundant Proteins in Experiments with Isobaric Labeling and Multiple Mixtures. Mol Cell Proteomics 19, 1706–1723. 10.1074/mcp.RA120.002105.

